# ERα-regulated IRX3 controls the growth of ER-positive breast tumors

**DOI:** 10.64898/2026.05.20.725898

**Authors:** Pouda Panahandeh Strømland, Jan-Inge Bjune, Regine Åsen Jersin, Mihaela Popa, Shuntaro Yamada, Kamal Mustafa, Emmet Mc Cormack, Karianne Fjeld, Elisabeth Wik, Simon Nitter Dankel, Gunnar Mellgren

## Abstract

Estrogen receptor positive (ER+) breast cancer is primarily treated with endocrine therapies targeting ERα signaling. Although endocrine therapy has substantially improved survival in ER+ breast cancer, metastatic disease remains largely incurable, underscoring the need to elucidate additional mechanisms driving growth and proliferation. Here, we show that the homeobox protein IRX3 is selectively overexpressed in ER+ breast cancer and define the molecular function of IRX3 in ER+ breast cancer using an integrated combination of *in vitro*, *in vivo* and *in silico* approaches. We uncover a previously uncharacterized distal regulatory region that controls *IRX3* transcription via ERα and associated steroid receptor coactivators. Consistent with this regulatory axis, anti-estrogen treatment resulted in marked downregulation of cellular IRX3 levels. Functionally, depletion of IRX3 suppresses proliferation of the human ER+ breast cancer cells *in vitro,* but paradoxically promotes tumor growth and metastatic dissemination in orthotopic xenografts *in vivo* by stimulating enhanced tumor vascularization. Finally, low tumor expression of IRX3 correlates with poorer survival outcomes in patients with ER+ breast cancer. Collectively, these findings establish IRX3 as an important regulator of ER+ breast tumor biology and reveal an ERα-dependent role for IRX3 in modulating proliferative and vascular programs in tumor progression.

**Significance:** By identifying a novel ERα-dependent regulatory pathway, this work refines our understanding of how hormone signaling shapes both breast tumor growth and the surrounding microenvironment.

## Introduction

Breast cancer is the most common form of cancer and the second-most common cause of cancer-related mortality in women worldwide (1). Breast tumors are highly heterogeneous, but 70-80% arise from luminal epithelial cells expressing high levels of estrogen receptor α (ERα) that drive tumor growth through increased cell proliferation and reduced apoptosis (2–4). ER signaling further supports tumor growth by increasing the tumor blood supply (5), either through mobilization and differentiation of endothelial progenitor cells into new blood vessels (neovascularization) or by enhancing growth of pre-existing capillaries through tubulogenesis and angiogenesis (6–9). The standard primary treatment for ER+ breast cancer is endocrine therapy, which reduces the ER signaling that drives tumor growth (10–12). Endocrine therapy commonly includes treatments that block estrogen action or reduce estrogen levels such as tamoxifen, fulvestrant and aromatase inhibitors (10–12). Emerging next-generation selective ER degraders, such as giredestrant, are also being developed to enhance endocrine therapeutic efficacy (12–15). However, many patients eventually develop resistance to endocrine therapy, and no curative options exist for advanced/metastatic disease (16). Thus, there is a need for new biological insights and therapeutic strategies.

To identify novel genes and mechanisms regulating growth and proliferation in ER+ breast cancer, we previously treated ER+ MCF-7 breast cancer cells with tamoxifen or its active metabolites and observed a marked reduction in mRNA expression of the Iroquois homeobox family protein 3 (IRX3), a highly conserved developmental transcription factor (17). Moreover, we have also previously reported that IRX3 promotes cell proliferation and cell-cycle progression in non-cancer preadipocytes *in vitro* (18). Other studies have reported that *IRX3* is derepressed in additional cancer types, including acute myeloid leukemia and hepatocellular carcinoma (19–21), and it is involved in the regulation of angiogenesis and vascularization (19–21). These findings suggest that IRX3 may be an ERα target gene, contributing in the regulation of ER+ breast cancer cell development and tumorigenesis.

In this study, we demonstrate that *IRX3* is selectively upregulated in ER+ breast tumors and is regulated by a previously unrecognized ERα-responsive enhancer. We further show that loss of IRX3 or targeted disruption of this enhancer, attenuates proliferative capacity in ER+ breast cancer cells *in vitro,* yet accelerates orthotopic tumor growth and metastasis *in vivo*. At the mechanistic level, repression of IRX3 elicits a compensatory proangiogenic and pro-vascular transcriptional program that facilitates tumor expansion despite reduced intrinsic proliferation.

## Methods

### Cell lines

The MCF-7, T-47D, and MDA-MB-231 cell lines were purchased from ATCC. The HEK 293T cell line, used to produce lentivirus particles, was kindly provided by Dr. J. Lorens. Human umbilical vein endothelial cells (HUVEC, C2517A, Lonza, USA) were a kind gift from Dr. Rolf Berg. MCF-7, MDA-MB-231 and HEK 293T cells were cultured in DMEM high glucose (4.5 g/L glucose, with glutamine and sodium pyruvate, VWR, Germany). T-47D cells were cultured in RPMI-1640 Medium. Media for MCF-7, T-47D and MD-MB-231 were supplemented with 10% fetal bovine serum, 1% (vol/vol) penicillin/streptomycin solution (Sigma, St. Louis, MO, USA). MCF-7 and T47-D cells media also contained 1 μM human recombinant insulin (Sigma). HUVEC were cultured in EGM2 medium with the manufacturer’s provided supplements (CC3162, Lonza, USA). All cells were cultured at 37°C in a humified incubator with 5% CO2 and atmospheric oxygen.

### E2-deprivation of the breast cancer cells

Breast cancer cells were seeded in 6-well plates and allowed to adhere overnight. The following day, cells were preconditioned for three days in phenol red-free DMEM (Invitrogen, Carlsbad, CA) containing charcoal-stripped fetal bovine serum (HycloneTM, Thermo Fischer Scientific, MA, USA) to deplete steroid hormones. After the E2-deprivation period, cells were treated for three days in the same pre-conditioned medium with or without 10 nM E2, either alone or in combination with 4-hydroxytamoxifen (4OH-tam, Sigma, Steinheim, Germany) or fulvestrant (Sigma).

### Generation of IRX3 knockdown cell lines

Two individual clones from MISSION® short hairpin RNA (shRNA) Target Set (Sigma) targeting human *IRX3* (shKD1: TRCN0000016899 and shK2: TRCN0000016902) and non-targeting scramble (shCtrl) were purchased. Each pLKO.1-puro-based vector was co-transfected with a lentivirus packaging plasmid into HEK 293T cells. Lentivirus particles were collected at 48 and 72 h after transfection, filtered and used to transduce cells in the presence of 8 μg/ml polybrene (Sigma). After 72-hr of puromycin (Sigma) selection, the infected cells were allowed to recover for at least 48 hours prior to testing for *IRX3* knockdown using real-time PCR.

### Plasmids, cloning and mutagenesis

A luciferase reporter plasmid containing the human *IRX3* promoter (pEZX-PG02-hIRX3p-1398/+290-GLuc) was purchased from Genecopoeia (Rockville, MD, USA). A 2kb intergenic region harbouring three ERα binding sites and exhibiting long-range interactions with the *IRX3* promoter was cloned into the pGL4.23-minP-empty luciferase reporter (Promega, Madison, WI, USA) to evaluate the enhancer activity of the genomic region. Two 5’ truncated versions of this enhancer (1.5 kb and 0.8 kb) were generated and cloned into the same vector. Genomic sequences were amplified by PCR from human genomic DNA using Platinum SuperFi Green PCR Master Mix (Invitrogen, Waltham, MA, USA) and primers pairs containing BglII or HindIII restriction enzyme (RE) sites at the 5’ end (**Supplementary Table S1**). The PCR products were purified by the QIAquick PCR purification kit (Qiagen, Hilden, Germany) and double-digested with BglII and HindIII-HF REs in NEBuffer 3.1 (New England Biolabs, Ipswich, MA, USA) alongside the empty vector. Digested inserts and vector backbone were purified by agarose gel extraction using the QIAquick gel extraction kit (Qiagen) before ligation using the Salt T4 DNA ligase kit (New England Biolabs) and a 3:1 insert : vector molar ratio. The ligation mixture was transformed into Top10 competent cells, and colonies were screened by colony PCR using vector and insert-specific primers (**Supplementary Table S1**). Positive clones were expanded and purified using the HiSpeed Plasmid Maxi kit (Qiagen) and verified by Sanger sequencing using BigDye 3.1 (Thermo Fisher, Waltham, MA, USA).

Site-specific mutagenesis of putative ERα response elements (EREs) was performed using the QuickChange II kit (Agilent, Santa Clara, CA, USA) according to manufacturer’s instructions except for the use of staggered primers (22) (**Supplementary Table S1**). All mutant constructs were confirmed by BigDye 3.1 Sanger sequencing.

The pSG5-empty and pSG5-ERα plasmids were kind gifts from Dr. E. Treuter. The pCDNA3-empty pCR3-SRC1, pCR3.1-empty, pCR3.1-SRC2 and pCR3.1-mSRC3 plasmids were kind gifts from Dr. Bert W. O’Malley. The pCAG-eCas9-GFP-U6-gRNA was a gift from Dr. Jizhong Zou (Addgene plasmid # 79145; http://n2t.net/addgene:79145; RRID: Addgene_79145).

### Genomic deletion of the IRX3 enhancer

To stably delete the enhancer, CRISPR-Cas9 genome editing was performed using two guide RNAs (gRNAs) targeting each flank of the enhancer followed by non-homologous end joining repair. The guides were screened using the UCSC genome browser track “CRISPR targets”, and the those with the highest predicted specificity and efficiency, that began with a G and were positioned as close as possible to the enhancer sequence were selected (**Supplementary Table S1**). The guides were cloned into the pCAG-eSpCas9-GFP-U6-gRNA-empty vector by Golden Gate assembly (23) using BbsI-HF (New England Biolabs) and validated by sequencing.

MCF-7 cells were transiently transfected with the two plasmids both containing eSpCas9-GFP and a guide RNA targeting either the 5’ or 3’ end of the enhancer. Briefly, 160,000 cells were seeded pr well in triplicates in a 6-well plate and transfected the following day using 1 µg of each plasmid and the XtremeGENE 9 DNA transfection reagent (Sigma Aldrich, St. Louis, MA, USA) at a DNA : reagent ratio of 1:5. Cells not receiving DNA were used as negative controls. Two days after transfection, GFP-positive cells were sorted using a Sony SH800S Cell Sorter, and 2,000 cells were subsequently plated in 10cm dishes to enable colony formation.

After 7-12 days, single colonies were picked under a microscope and transferred to a 96-well plate. After ≥14 days, proliferating colonies were expanded to 24-well plates for further expansion and genomic DNA (gDNA) extraction. For the initial screening, gDNA was extracted using the GeneElute mammalian genomic DNA miniprep kit (Sigma). Clones were screened by qPCR using primers binding either inside, outside or spanning both sides of the enhancer to identify successfully edited clones (**Supplementary Table S1**). One of the clones with minimal signal inside the enhancer was subjected to a second round of colony formation and 96 subclones were screened using Chelex-based gDNA extraction and qPCR screening. Identification of pure clones with complete loss of the enhancer was validated by agarose gel electrophoresis and sanger sequencing.

### Luciferase assays

On day 1, a total of 8,000 MCF-7 or MDA-MB-231 cells were seeded per well in a 96-well plate in E2-deprived medium and allowed to grow overnight (o/n). The next day, the cells were given fresh medium and transfected with 30 ng of reporter plasmid and 40 ng of either empty plasmid, ERα and/or SRC1, SRC2 or SRC3 per well, as indicated. After 24h, the medium was replaced with fresh E2- deprived medium supplemented with either DMSO, 10 nM E2, 250 nm 4OH-tam or 100 nm fulvestrant. The following day, the cell culture media were harvested for measurement of secreted Gaussia luciferase (promoter constructs) or the cells were lysed (in 25 mM TAE pH 7.8, 1% Triton-X, 1 mM EDTA, 2 mM DTT, and 10 % glycerol) for measurement of non-secreted Firefly luciferase (enhancer constructs). Gaussia and firefly luciferase activity was measured using the Pierce Gaussia Luciferase Glow Assay Kit (Thermo Fisher) and Luciferase Assay Kit (Biothema, Handen, Sweden), respectively, according to manufacturers’ instructions. Raw luciferase units were normalized to the average of the control samples to obtain relative luciferase units.

### Real-time qPCR

Cells were lysed in RLT buffer and RNA was extracted using the RNeasy mini kit (Qiagen) followed by cDNA synthesis using 1,000 ng input and the High-Capacity cDNA Reverse Transcription Kit (Thermo Fisher). The cDNA was diluted 1:5 in RNase-free H_2_O and qPCR was performed using the LightCycler® 480 SYBR Green I Master (Roche, Basel, Switzerland) with 1.25 µL template and 40 pmol of each primer in a total volume of 10 µL. The relative amount of cDNA was calculated by the ΔΔCtCT method with *PUM1* and/or *TBP* as reference genes. In experiments where cells were pre-conditioned with E2, expression of *GREB1,* an early ER-responsive gene, was used as positive control. All experimental conditions were performed with a minimum of three biological and three technical replicates. Primer sequences are listed in (**Supplementary Table S1**).

### Subcellular fractionation

Control and Δenhancer MCF-7 cells grown in 10 cm dishes were harvested by scraping, centrifugated at 300 x g for 3 min, and the resulting pellets were snap frozen in liquid nitrogen before storage at −80°C until fractionation. On the day of fractionation, cells were thawed on ice and resuspended in 500 µL hypotonic buffer (10 mM HEPES pH 7.8, 1.5 mM MgCl_2_, 10 mM KCl, 0.1% IGEPAL, 0.5 mM DTT) supplemented with 1X cOmplete Ultra protease inhibitor and 1X PhosSTOP (both from Roche), followed by swelling for 30 min on a spinning wheel at 4°C and gentle sonication (1-3X 15 sec on/30 sec off on LOW setting on a Nexon sonicator). Release of the nuclei was monitored by Trypan blue staining and brightfield microscopy. The nuclei were pelleted at 13,000 x g for 5 min at 4°C and the supernatant containing cytosolic proteins was removed and saved for subsequent analyses. The pellet was washed 1X in the same buffer before being thoroughly resuspended in 100 µL of a high-salt buffer (20 mM HEPES pH 7.8, 1.5 mM MgCl_2_, 420 mM NaCl, 0.2 mM EDTA and 0.5 mM DTT supplemented with 1X cOmplete Ultra protease inhibitor (Roche) and 1X PhosSTOP). The suspension was subjected to moderate sonication (≥3X 30/30 sec on MEDIUM settings on a Nexon sonicator). The lysate was then centrifugated at 13,000 x g for 15 min at 4°C, and the supernatant containing nuclear proteins was collected for subsequent analyses.

### Immuno-blotting

Cell lysates were quantified using the DC Protein Assay (Bio-Rad, Hercules, CA, USA) and 20 µg normalized protein of each sample was analyzed by standard western blotting. Primary antibodies used were anti-IRX3 (Cell Signaling, 28398), anti-p57 (Cell Signaling, 2557), anti-CDK2(D-12, Santa Cruz, Dallas, TX, USA, sc-6248), anti-H3 (Cell Signaling 3638S) and secondary antibodies were anti-rabbit (Thermo Scientific, 31460) and anti-mouse (BD Biosciences, 554002).

### Proliferation and 2D cell cycle assays

In the proliferation assay, a total of 2,000 cells (n≥6 for each condition) were seeded per well in a 96-well plate. Cell growth was monitored by high content imaging using the IncuCyte S3 Live-Cell system every 4 h for 3-5 days. Images were analyzed using the IncuCyte S3 software, and cell growth is presented as percentage of confluency normalized to *t* = 0. For the 2D cell cycle assay, 250,000 non-synchronized cells were grown in 6-well plates for 48 h before being pulse-labeling for 2 hours with 30 µM BrdU (Abcam). The cells were then incubated overnight in BrdU-free medium, fixated in 70% ethanol, permeabilized and stained with anti-BrdU (Abcam, ab6326) or negative control IgG (Abcam, ab171870) as negative control and Propidium Iodide (PI). The stained cells were analyzed by flow cytometry using Accuri C6 (BD Biosciences) and data were analyzed with FlowJo software (Tree Star, Inc) as previously described (17).

### Endothelial cell proliferation assay using conditioned media

A total of 1,500 cells (n=5 for each condition) of human umbilical vein endothelial cells (HUVEC) were seeded per well in a 96-well plate. Conditioned media from 48 h of cultured IRX3-knock down MCF-7 cells was mixed 80:20 (v/v) with fresh HUVEC medium and added to the HUVEC cells the day after. The growth of HUVEC cells was monitored for 4 days as other proliferation assays.

### *In vivo* experiments

Mice experiments were approved by the local Ethical Committee at the University of Bergen and the Norwegian Food and Safety Authority (application ID 28545) in accordance with the Norwegian Commission for Laboratory Animals.

### Immunodeficient mice

Female NOD. Cg-Prkdcscid Il2rgtm1Wjl/SzJ (6-10 weeks old; Strain #005557) mice were purchased from Charles River Laboratories. Mice were housed in individually ventilated (HEPA-filtered) cages (Techniplast, Buguggiate, Italy) under defined flora conditions. Cages were kept on a 12-hour dark/light schedule at a constant temperature of 21°C and 50% relative humidity. Mice had continuous access to sterile water and food and were monitored daily by the same staff throughout the experimental period.

### Xenograft models

For xenograft ER+ cell models, the established MCF-7 IRX3-KD and IRX3 Δenhancer cells were transduced with 10 MOI RediFect Red-FLuc-GFP Lentiviral Particles (PerkinElmer, Product No. CLS960003). After expansion under normal conditions, the cells with the highest GFP signals were sorted using the Sony SH800S Cell Sorter. The sorted GFP+ cells were then expanded, and the phenotype was validated (preservation of reduced *IRX3* expression and growth rate of the GFP+ shKD and Δenhancer cells). Before cell implantation, luciferase expression was verified by adding 10 µl D-luciferin (Promega, Madison, WI, USA, 150 µg/µl) to 100 µl cell suspensions containing 100,000 cells in a 96-well plate, followed by optical imaging.

The tumor growth and metastatic potential of the IRX3 knock-down and IRX3 Δenhancer cells were evaluated and compared to the corresponding luciferin expressing MCF-7 control cells. The cells were implanted orthotopically into the inguinal mammary fat pad of the NSG mice. For each cell line, mice (n = 8 per group) were injected with 1,000,000 cancer cells resuspended in 50 µL sterile saline mixed 3:1 with Matrigel (Corning, Cat. No. 354248). MCF-7 cells were implanted into mice receiving continuous E2 supplementation from slow-release rods (PreclinApps) implanted 48 h prior to cancer cell injection. The rods delivered a sustained release of 1.5 µg E2 per day for approximately 100 days, supporting tumor growth.

### Optical imaging

Weekly optical imaging of all the mice in the study was performed using the IVIS Spectrum system (Revvity). Prior to imaging, mice with engrafted tumors received a contrast reagent, D-Luciferin (150 mg/kg, 250 mg/mL), via intraperitoneal injection. The BLI intensity was quantified for each mouse, and the values were plotted for each experimental group to monitor and analyze disease progression over time. *Ex vivo* images of organs with metastases were acquired at the end of the experiment to assess and compare the extent of metastatic spread between mice engrafted with IRX3-KD cells and those engrafted with IRX3 Δenhancer cells. Data was analyzed using Living Image 4.7.3. software.

### Histological evaluation and immunohistochemical staining of FFPE tumor sections

Tumor tissues from animal experiments were collected at the endpoint. Primary tumors were divided into two: one part was snap-frozen for RNA extraction and sequencing (see method below), and one part was fixed in 4% formalin in PBS for histological staining. For the latter, mice presenting with the lowest IRX3 expression in primary tumor (assessed by RT-PCR) and pronounced in vivo phenotype (based on tumor weight) were analyzed (n=4). Formalin-fixed and paraffin-embedded (FFPE) tumor tissue was sectioned (4-5 mm) and stained using standard protocols at the Department of Pathology, Haukeland University Hospital.

Hematoxylin and Eosin (HE) and Masson Trichrome staining (Roche, 860-031) were used to evaluate histology and fibrosis development, respectively. A monoclonal mouse anti-human Ki67 antibody (Dako, M7240) was used as a marker for cell proliferation. Following staining, the tissue sections were cover-slipped, and whole-slide scanning (40x) was performed using an Aperio AT2 (Leica Biosystems).

Mitotic count and necrosis were performed by an experienced pathologist (E. W.) on randomized, anonymous HE stained tumor sections. To quantify the mitotic cells in each tumor section, the most cellular and the least necrotic area with the highest number of mitotic figures (hot spot) was marked for counting. Mitotic figures were counted in 10 consecutive high-power fields (x400) and reported per mm². Caution was taken for areas with fibrosis, and the presence of immune cells. The percentage of tumor necrosis was defined as the proportion of areas of necrotic tumor cells relative to the total tumor area (mm2). Single-cell necrosis between the viable parts of the tumor was not included in the necrotic area. Quantification of Ki67 and Masson’s trichrome staining was performed using QuPath (version 0.7.0) (24).

### *Ex ovo* chorioallantoic membrane (CAM) assay

Fresh fertilized eggs of *Gallus gallus domesticus* were incubated at 37.5 °C for 3 days in humidified conditions using a rotating egg incubator. The egg contents, including chicken embryos, were subsequently transferred into sterile weighing boats with 2 ml of PBS and incubated at 37.5 °C for an additional 4 days. On embryonic day 7, 3 cell-laden Matrgels, prepared with 200,000 MCF-7 cells from each condition, were placed onto each chorioallantoic membrane (CAM) within a Teflon ring. On embryonic day 12, the CAM was dissected and fixed in 4% PFA overnight to enable blood vessel quantification. For quantification, the microscopic images were obtained using a stereomicroscope (M205C/MC170HD, Leica, Germany) and quantified using a deep learning-based image analysis software, IKOSA CAM Assay Application (KML Vision GmbH, Austria).

### Transcriptomics analyses

#### Human clinical data

Total transcriptomics profiles of paired normal- and invasive tumor breast tissues from patients with carcinoma were obtained from unpublished data (GSE70947, n = 147). Differentially expressed genes were identified using the limma Bioconductor package in R (25), applying adjusted p-value cut-off of 0.01.

### Single cell RNA sequencing data

The processed scRNAseq data was obtained from Gene Expression Omnibus GSE161529. The analysis of the data was replicated as described previously (26). Data from samples from each normal and tumor subtypes were combined using an anchored-based integration method using Seurat package in R (27). The cell clusters were identified using the Louvian clustering algorithm in Seurat. Principal component dimensions 1:30 and clustering resolutions 0.1 were used for all dimensions reduction and integration steps. The epithelial cell cluster was defined by *EPCAM* expression in all normal and tumor breast tissues, also confirmed by the expression of myoepithelial, luminal progenitor- and mature luminal cell markers.

### RNA sequencing

For transcriptomic analyses, cells were lyzed for RNA extraction as for real-time PCR, with an additional on-column DNase-I treatment (Qiagen, 79254) performed at 27 °C. RNA purity and integrity (RIN) were quantified using the RNA 6000 Nano kit (Agilent Technologies, 5067-1511) on the 4200 TapeStation (Agilent, Santa Clara, USA). Library preparation was performed from 160 ng total RNA using the Illumina Stranded Total RNA Prep Ligation kit, and sequencing (2×75bp paired end runs) was carried out on the Illumina HiSeq4000 system (Illumina, Sand Diego, CA, USA. Raw reads from the HiSeq4000 were quality checked and aligned to the human reference genome GRCh38 (GCA_000001405.15) using HISAT2 2.0.5 and submitted to Subread v.1.5.2 for feature counts calculation.

Transcriptomic analyses were performed on *in vitro* GFP+ MCF-7 IRX3-KD and Δenhancer cells (n = 4 each group) and snap-frozen murine primary tumors obtained from xenografts of shCtrl, shKD2, Ctrl, and ΔEnh1 cells (n = 5 each group), selected based on *IRX3* expression performed by real-time PCR). RNA from snap-frozen cells and primary tumors were purified after being lyzed using Tissuelyser (Qiagen) and the RNeasy Mini kit (Qiagen, 74116) with additional on-column DNase-I treatment (Qiagen, 79254) performed at 25 °C.

RNA purity and integrity (RIN) were measured using the RNA 6000 Nano kit (Agilent Technologies, 5067-1511) on the 4200 TapeStation (Agilent, Santa Clara, USA). Complementary DNA (cDNA) libraries were prepared using the Illumina Stranded mRNA Ligation kit with 350ng of input RNA. All samples were quality-controlled by the 4200 TapeStation using the DNA ScreenTape Analysis. Final cDNA libraries were additionally quantified with the KAPA Library Quantification Kit (Roche, USA) for Illumina sequencing platforms. All samples were paired end sequenced (2×100bp) on the Illumina NovaSeq 6000 system (Illumina, San Diego, CA, USA). Run data from NovaSeq6000 were demultiplexed by the Illumina bcl2fastq software, and the Fastq files were quality-controlled by FastQC and MultiQC. Reads were aligned to the human reference genome GRCh38 (GCA_000001405.15) and transcriptome GENCODE_vM13 reference using HISAT2 2.2.1 and subsequently submitted to Subread v.1.5.2 for feature counts calculation, resulting in a raw read count matrix.

The differentially expressed genes was identified using DESeq2 Bioconductor package in R (28). Gene ontology (GO) enrichment analyses of the differential expressed genes (adjusted p-value <0.01) between the IRX3-KDs and shCtrl, and Δenhancer and control were performed using PANTHER 19.0 (29). Significant enriched GO terms were selected as FDR < 0.05 and GOs with overrepresentation of genes were depicted (**Supplementary Dataset 1-2**). Gene expression signature scores were calculated for VEGF gene signatures that are associated with vascularization and angiogenesis (30). Mean z-score of all genes in the VEGF signature set constituted the score for this signature.

### ChIA-PET and ChIP sequencing analyses

For analysis of long-range chromatin interaction at the *IRX3* locus and putative cis-regulatory regions on chromosome 16, publicly available RNA polymerase II (RNA POLII) Chromatin Interaction Analysis by Paired-End Tag Sequencing (ChIA-PET) data from MCF-7 cells were obtained from the Washington University Epigenome Browser (http://epigenomegateway.wustl.edu/browser/). All the ChIPseq data obtained from published studies were downloaded from Gene Expression Omnibus. To obtain the precise coordinates corresponding to the *IRX3* promoter and the predicted enhancer, the raw data were downloaded and aligned to the human genome GRCh38.

### Relapse-free survival analysis

Relapse-free survival (RFS) was assessed using the KMplotter tool (31) in an integrated dataset of three cohorts (The cancer genome atlas (TCGA, https://www.cancer.gov/tcga), METABRIC (32) and IMPACT (33)) including 2,032 breast cancer patients. Median IRX3 expression level was used to stratify patients within each tumor subtype into high- and low-expressing groups.

### Statistical analyses

Raw RNA sequencing data were analyzed in R as described above. The remaining data were analyzed and graphed using GraphPad Prism (9.2.0). Statistically significant differences between group means were investigated using two-sided Student’s unpaired t-test, one-way ANOVA, or two-way ANOVA with Holm-Sidak correction for multiple testing as indicated. A two-sided adjusted *p*-value < 0.05 was considered statistically significant. Data were tested for normality and homogeneity of variance, and transformed, if necessary, prior to statistical analysis, to conform with the assumptions of the tests. In animal experiments, outliers, when caused by obvious technical errors, such as loss of the E2 rod, were removed. The Kruskal–Wallis test was used to compare non-parametric and continuous variables across groups for histological parameters. For univariate relapse-free survival analyses, Kaplan–Meier product-limit method (log-rank test) was used in KMplotter tool.

### Data availability

Global gene expression data derived from the IRX3-KD1, and 4OH-tamoxifen-treated MCF-7 cells have been deposited in Gene omnibus (GEO) database (GSE301355). The transcriptomics data derived from the IRX3-KDs and IRX3 Δenhancer GFP+ MCF-7 cells as well as from the primary tumor-xenografts in mice are accessible at GEO database (GSE301481).

## Results

### IRX3 is overexpressed in estrogen-receptor positive epithelial breast cancer cells

To investigate the potential role of IRX3 in ER+ breast cancer, we interrogated a clinical dataset (GSE70947) to compare the expression levels of *IRX3* in 148 paired normal and tumor breast tissue samples from both ER- and ER+ breast cancer patients (**Figure 1A**). ER+ tumors subtype showed higher expression of *IRX3* compared to both matched normal breast tissue (∼3-fold change, *p_adj_* < 0.01) and ER- tumor tissues (∼2-fold change, *p_adj_* < 0.01). *IRX3* expression was also significantly and positively correlated with *ESR1*, the gene encoding ERα (**Figure 1B**). The single-cell RNA-sequencing (scRNA-seq) atlas (26) confirmed increased *IRX3* expression in ER+ breast cancer compared to other subtypes, with elevated *IRX3* expression specifically in cell clusters expressing high levels of *ESR1* (**Figure 1C**). Moreover, the expression pattern of *IRX3* closely paralleled that of *EPCAM*, a key epithelial cell marker, across diverse cell populations in both breast and tumor subtypes, suggesting that *IRX3* expression is confined to epithelial compartments in both normal and tumor tissues (**Figure 1C**). Identification of the three major epithelial cell clusters in normal breast tissue further demonstrated that *IRX3* is more highly expressed in luminal progenitor and mature luminal cells than in the basal/myoepithelial compartment (Cluster 0 and 2, **Supplementary Fig. S1A**). Taken together, these findings demonstrate that *IRX3* expression is positively correlated with *ESR1* expression in breast tissues.

**Figure 1.**
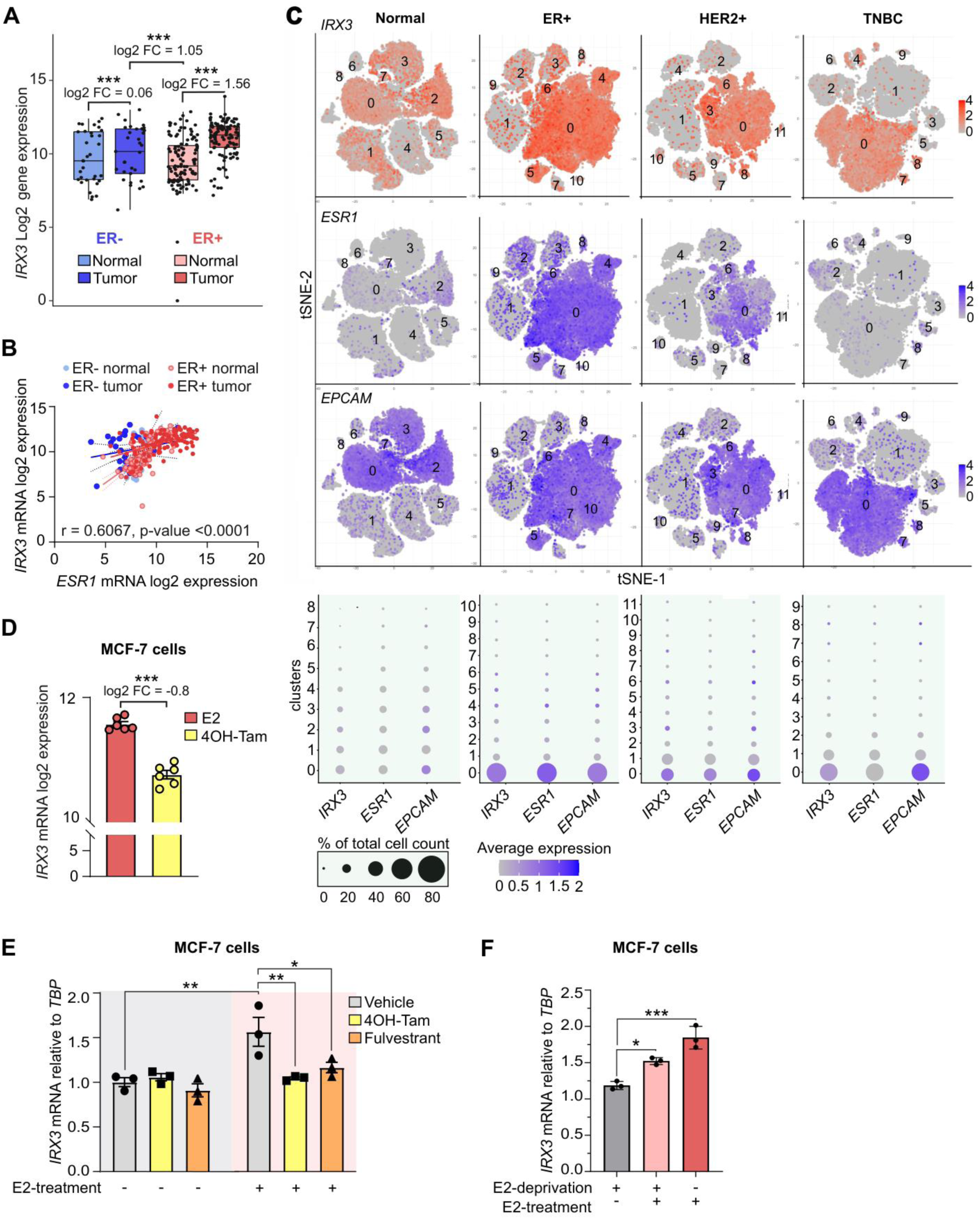
*IRX3* displays elevated expression in estrogen receptor positive epithelial breast cancer cells. **A.** The mRNA expression levels of *IRX3* in ER- and ER- paired normal and tumor breast tissues (n = 148 breast cancer patients). The data are derived from microarray RNA expression profiling from an unpublished clinical study (GSE70947). Log2 fold changes (FC) from the differential gene expression analysis (adjusted p-value = 2.67e^-06^) are presented. \*\*\**p_adj_* < 0.001. **B.** The *Estrogen Receptor 1 (ESR1)* and *IRX3* log2 transcript expression correlation in ER- and ER+ paired normal and tumor breast tissues (n = 148) shown in A. Spearman’s correlation coefficient and corresponding p-values are shown. **C.** The t-distributed Stochastic Neighbor Embedding (tSNE) plots show *IRX3*, *ESR1* and *EPCAM* expression in normal breast tissue (n = 13), as well as in ER+ (n = 17), HER2+ (n = 6) and Triple-Negative Breast Cancer (TNBC; n = 8) breast tumor tissue. The data is derived from single-cell RNA sequencing (scRNAseq) (26) (GSE161529). The percentage of cells within each cluster and average expression level (normalized transcript count) is shown in the dot plot. **D.** Expression levels of *IRX3* in MCF-7 cells. Cells were exposed to E2-deprived medium for 72 h, and subsequently treated with 10 nM estradiol (E2) with or without 250 nM 4-OH-tamoxifen (4OH-tam) for 48 h. The data are derived from a microarray mRNA expression dataset (n = 6 for each treatment (16) (E-MTAB-2729). Bar graphs show means ± SEM. Log2 fold changes (FC) from the differential gene expression analysis (adjusted p-value = 1.8e^-08^ is presented) are presented. \*\*\**p_adj_* < 0.001. **E.** Expression levels of *IRX*3 in MCF-7 cells during treatment with 4OH-tam and fulvestrant. MCF-7 cells were cultured in E2-deprived media for 72 h and subsequently treated or not treated with 10 nM estradiol (E2), with or without 4OH-tam (250 nM) or fulvestrant (100 nM) for 48 h. Real-time PCR of *IRX3* relative to the TATA-Binding Protein (*TBP*) house-keeping gene was performed. The data are derived from n = 3 biological replicates and representative of two independent experiments. \**p_adj_* < 0.05, \*\*\**p_adj_* < 0.001; Two-Way Anova with Holm-Sidak correction for multiple testing. Bar graphs show means ± SD. **F.** Restimulation of *IRX3* expression in E2-deprived MCF-7 cells. MCF-7 cells were pre-conditioned in E2-deprivation medium with or without 10 nM E2 for 72 hr. The E2-deprived cells were then treated or not treated with 10 nM E2 for 72h. All cells were harvested at the same time. *IRX3* mRNA expression was quantified by real-time PCR relative to *TBP* house-keeping gene expression. The data are derived from n = 3 biological replicates. \**p_adj_* < 0.05, \*\**p_adj_* < 0.01, \*\*\**p_adj_* < 0.001; Two-Way Anova with Holm-Sidak correction for multiple testing. Bar graphs show means ± SD.

To investigate whether *IRX3* expression is controlled by ERα signaling, we first analyzed transcriptome profiles of MCF-7 ER+ breast cancer cell line treated with the active tamoxifen metabolite 4-hydroxytamoxifen (4OH-tam) in a previous *in vitro* study (16), and confirmed *IRX3* downregulation by ∼40% reduction (**Figure 1D**). Notably, *IRX3* was among the top 20 downregulated genes in this experiment (**Supplementary Fig. S1B**). Moreover, upon silencing *IRX3* expression in MCF-7 cells, we found a striking 71% overlap of differentially expressed genes in the same direction as upon 4OH-tam treatment (152 out of 213 commonly differentially expressed genes) (**Supplementary Fig. S1C**). Consistently, when restimulating 17β-estradiol (E2)-deprived MCF-7 cells with E2, we observed a significant increase in *IRX3* expression, and a reversal of this effect by both 4OH-tam and the ER antagonist fulvestrant (**Figure 1E**). Additionally, after culturing MCF-7 cells in E2-deprived medium for three days, reintroduction of E2 significantly increased *IRX3* expression towards the level of non-deprived cells (**Figure 1F**). These data show that *IRX3* is a target gene of ERα and mediates a substantial part of the transcriptional response to 4OH-tam.

### ERα controls *IRX3* expression via an upstream enhancer

To further investigate the regulation of *IRX3* by estrogen-ERα signaling, we searched for conserved ERα response elements (EREs) in the proximal *IRX3* promoter. JASPAR (34) enrichment analyses predicted a putative ERE of about 600 bp upstream of the transcription start site (**Supplementary Fig. S2A**). Therefore, we co-expressed a luciferase reporter harboring the *IRX3* promoter together with an expression plasmid encoding ERα and tested for transcriptional regulation of *IRX3* by E2 or 4OH-tam in E2-deprived ER+ MCF-7 as well as ER- MDA-MB-231 cells (**Supplementary Fig. S2B**). However, neither E2 nor 4OH-tam affected reporter activity (**Supplementary Fig. S2B**). We also interrogated publicly available ERα ChIP-seq studies in ER+ tumors and MCF-7 cells (35,36), which consistently showed no significant ERα binding to the promoter of *IRX3* (**Supplementary Fig. S2C**).

We next hypothesized that ERα regulates *IRX3* expression via one or more enhancers rather than the promoter. We and others have previously shown that *IRX3* expression is regulated by a distal enhancer located in intron 1 of the *FTO* gene in human adipose-derived mesenchymal stem cells (37–39). To look for distal chromatin regions in contact with the *IRX3* promoter in ER+ breast cancer cells, we searched for publicly available long-range chromatin interaction data (see Methods section) and found strong interactions between the *IRX3* promoter, and an intergenic region located approximately 140Kb upstream, as well as intronic regions in *FTO* and the promoter of *IRX5* (**Figure 2A**, top panel). ERα ChIP-seq data from MCF-7 cells and primary tumors of ER- and ER+ breast cancer patients (35,36) also showed strong and consistent binding of ERα to the same location (**Figure 2A**, bottom panel). Closer inspection of the ERα binding signal at this region revealed three distinct peaks within a 2 kb window, which overlapped with conserved EREs as predicted by JASPAR (**Figure 2B, C**). Open chromatin and epigenetic data indicate that these ERα binding sites, located in the predicted enhancer may be active in breast tissue (**Figure 2B**).

**Figure 2.**
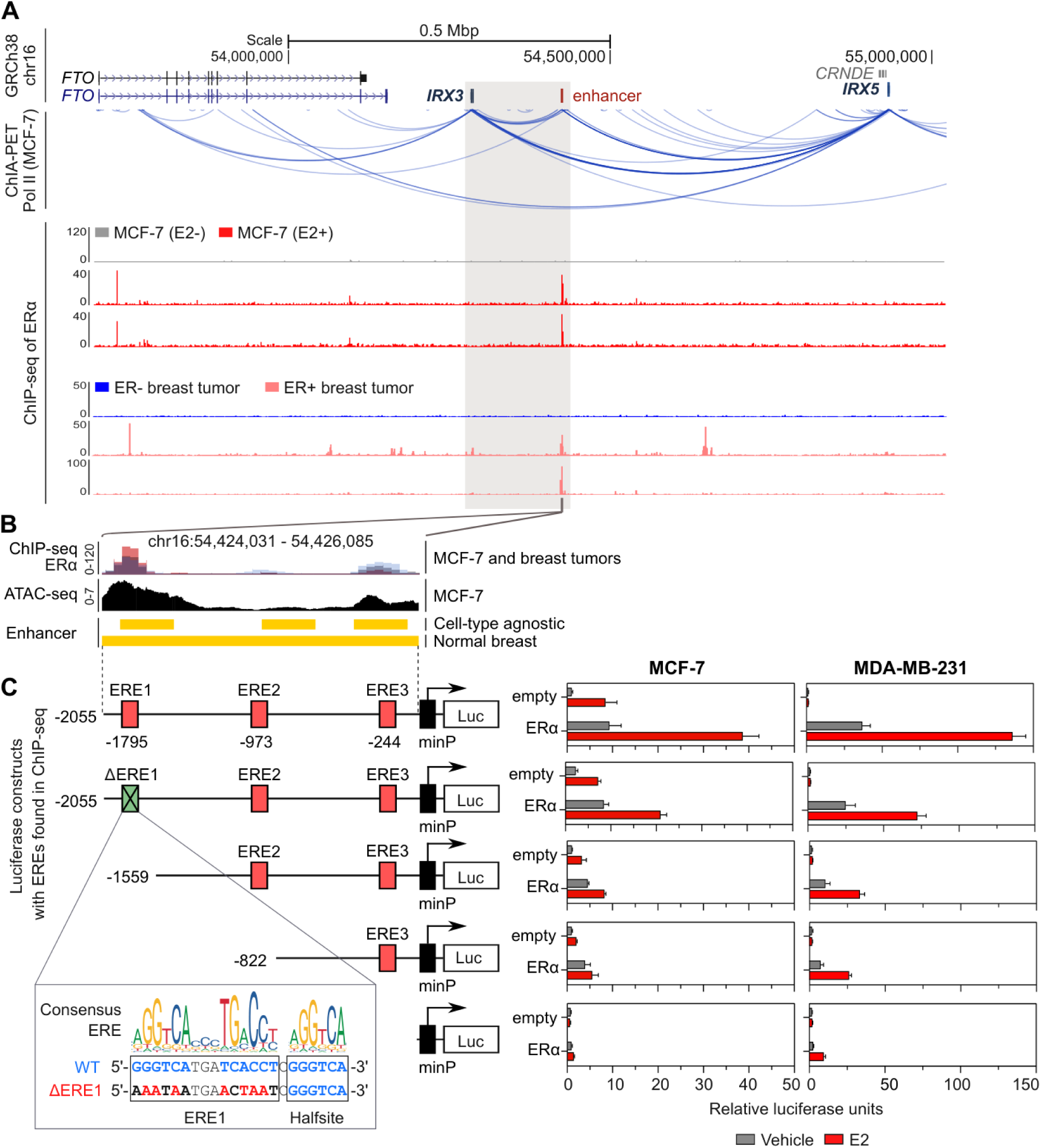
ERα controls *IRX3* expression via an upstream enhancer. **A.** Genomic view of the *FTO*-*IRX3* locus (top panel). Chromatin Interaction Analysis by Paired-End Tag Sequencing (ChIA-PET) map of RNA POL II (Pol II), with arcs showing interactions between any two regions in chromosome 16 of MCF-7 cell genome (middle panel). Stronger color denotes stronger interaction. The data is derived from http://epigenomegateway.wustl.edu/browser/ (see Methods section). ChIP-seq tracks (lower panel) showing ERα binding in the presence (E2+) or absence (E2-) of 10 nM 17β-estradiol in MCF-7 cells and ER- and ER+ primary breast tumors derived from breast cancer patients. The grey box highlights chromatin interactions between *IRX3* and a nearby locus with ERα ChIP-seq peaks (*IRX3* enhancer). The data is derived from published studies GSE32222 and GSE14664 (35,36). **B.** Zoomed-in view of the *IRX3* enhancer. Overlaid ERα ChIP-seq tracks from E2+ treated MCF-7 cells and ER+ tumors from Fig. 2A (top panel), open chromatin in MCF-7 cells (middle panel), and ENCODE/Roadmap epigenetic enhancer marks (bottom panel) shown. The data is derived from http://epigenomegateway.wustl.edu/browser/ and https://genome.ucsc.edu/. **C.** Schematic overview of the wild-type (WT) enhancer cloned into the pGL4.23-minP-empty luciferase vector (left panel). JASPAR-predicted EREs are shown as red boxes. WT full-length enhancer, site-directed mutation of the most distal ERE in the full-length enhancer (ΔERE1, green box) and 5’ deletion constructs are shown. Relative luciferase activity of each reporter construct with and without co-expression of ERα in the presence or absence of E2 in E2-deprived ER+ MCF-7 and ER- MDA-MB-231 cells (right panel). Data are derived from n = 4 biological replicates and representative of three independent experiments. Bar graphs show means ± SD. The consensus ERE shown as position weight matrix logo, aligned with the genomic sequence of WT and mutant ERE1 (ΔERE1). ERE, ERα response element; Luc, luciferase reporter gene; minP, minimal promoter.

Based on the hypothesis that this 2 kb locus, located approximately 140 kb upstream of *IRX3* could act as an ERα-responsive enhancer controlling *IRX3* expression, we cloned the region into a luciferase reporter construct under control of a minimal promoter (pGl4.23-minP-IRX3enh-luc) (**Figure 2C**, left panel). The reporter was co-expressed with ERα in the presence or absence of E2 in E2-deprived MCF-7 and MDA-MB-231 cells. While the basal enhancer activity was low, ERα overexpression and addition of E2 increased the enhancer activity 40-fold in E2-deprived MCF-7 cells (**Figure 2C**, right panel). In the ER- MDA-MB-231 breast cancer cell line, the enhancer activity increased nearly 150-fold by ERα overexpression and addition of E2 (**Figure 2C**). Site-directed mutagenesis of the ERE with the strongest ERα ChIP-seq signal (ΔERE1), as well as an accompanying predicted half-site (ΔERE5) reduced the enhancer activity by 50% and 80%, respectively (**Figure 2C, Supplementary Fig. S3**), demonstrating that the majority of ERα binding occurred at this genomic region. Additional deletion constructs and site-directed mutants, including several additional predicted EREs and half-sites, further revealed that ERE3 contributes to enhancer activity, with minor contributions from multiple additional half-sites (**Figure 2C, Supplementary Fig. S3**). Taken together, the 2 kb locus upstream of *IRX3* physically interacts with the *IRX3* promoter and acts as a strong ERα-responsive enhancer in ER+ breast cancer cells *in vitro*.

ERα-mediated transcriptional activity is regulated by interactions with the steroid receptor coactivators (SRC) 1, −2 and −3 (40). A publicly available ChIP-seq dataset (40) (E-MTAB-785) showed that SRC-2 and SRC-3 binding in the MCF-7 genome particularly overlapped with ERα binding at the enhancer region (**Supplementary Fig. S4A)**. We therefore overexpressed the SRCs together with ERα in ER- MDA-MB-231 cells and assessed their ability to activate the pGl4.23-minP-IRX3enh-luc luciferase reporter in the presence of E2. SRC-1 and SRC-2 demonstrated minimal effects on the enhancer alone but greatly increased the E2/ERα-dependent enhancer activity (**Supplementary Fig. S4B**). SRC-3 appeared to affect the enhancer independently of ERα as previously reported for some cis-regulatory regions (40,41). Moreover, mutation of the ERE1 (ΔERE1) strongly reduced the enhancer activity imposed by ERα alone and in synergy with the SRCs, while not affecting the ERα-independent effect of SRC-3 (**Figure S4B**). We next examined whether 4-OH-Tam and fulvestrant could impair the ERα/SRCs-dependent activation of the enhancer. Both anti-estrogens markedly inhibited ERα/SRCs-mediated enhancer activity in the presence of E2 in both MCF-7 and MDA-MB-231 cells (**Supplementary Fig. S4C**). Overall, these data further support the enhancer to be activated by estrogen-bound ERα and coactivators.

To experimentally validate that *IRX3* is the target gene of the ERα-responsive enhancer, we used CRISPR-Cas9 to delete the 2 Kb enhancer region in the genome of MCF-7 cells (**Figure 3A**). Strikingly, deletion of the enhancer reduced *IRX3* mRNA levels by 60-80% in three different clones (**Figure 3B**). In contrast, *IRX5*, whose promoter also showed chromatin interaction with the enhancer, was only weakly affected in one of the Δenhancer clones (**Figures 3B**), suggesting that *IRX3* may be the primary target of the enhancer. Importantly, IRX3 protein levels decreased by 80-90% in two clones (Δenh1 and Δenh3) (**Figure 3C**). Taken together, these data demonstrate that the ERα-responsive enhancer regulates *IRX3* expression in MCF-7 cells and plays a critical role in maintaining IRX3 levels.

**Figure 3.**
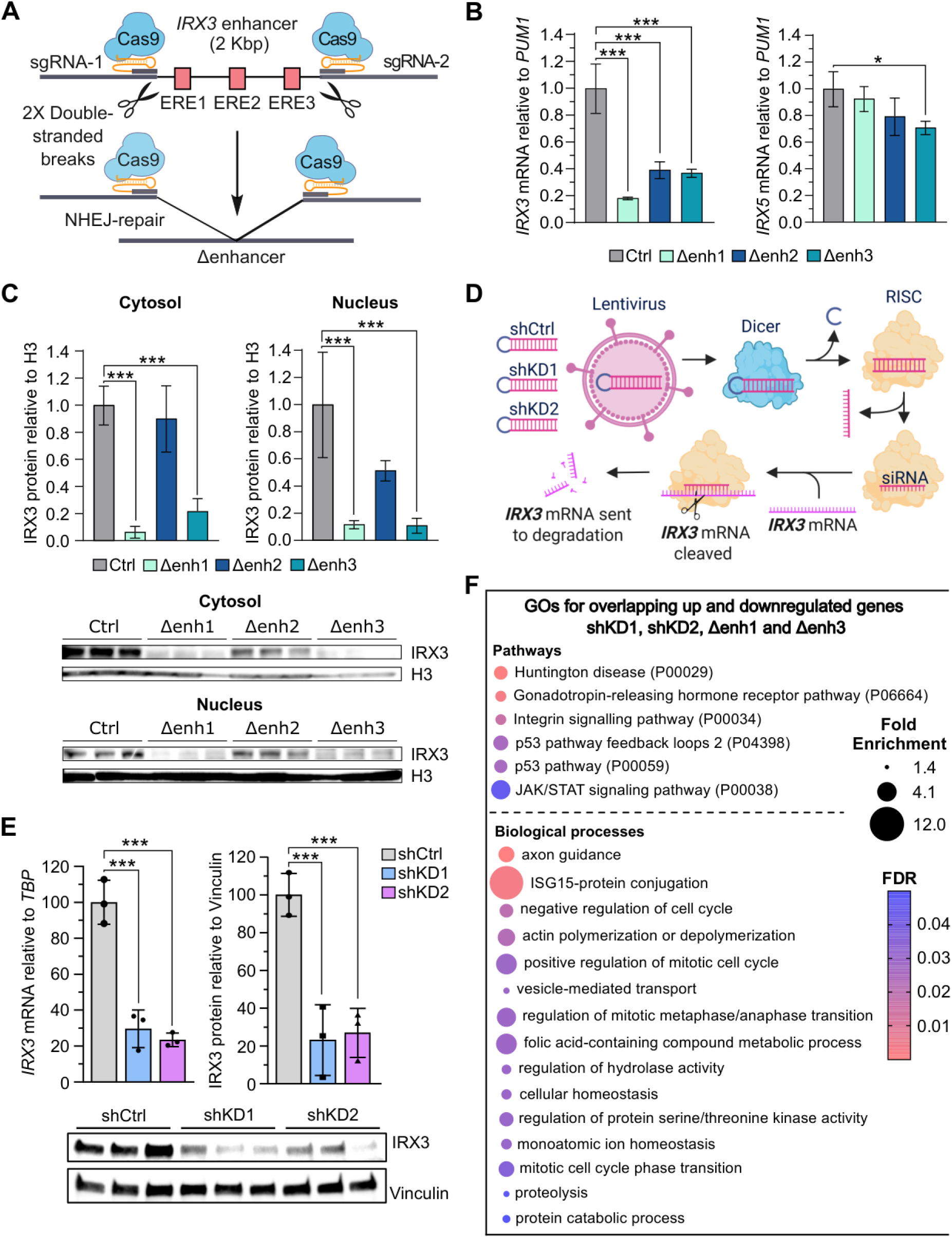
Decreased IRX3 levels by genomic deletion of the *IRX3* enhancer or direct targeting of *IRX3* transcripts by short hairpin RNA affect expression of genes involved in proliferation and regulation of the cell cycle. **A.** Schematic illustration of the strategy for deletion of the IRX3 enhancer in MCF-7 cells using CRISPR-Cas9. Two sgRNAs were designed to flank each site of the enhancer. Following transfection of plasmids encoding the sgRNAs and Cas9-T2A-GFP, GFP+ cells were sorted by FACS and colonies from single cells expanded. Clones with successful double-stranded breaks and non-homologous end joining (NHEJ) repair that excised the enhancer were identified by qPCR and validated by Sanger sequencing. Clones transfected with Cas9, but not sgRNAs, were used as control. The figure is created in BioRender. **B.** The expression of *IRX3*, *IRX5* relative to housekeeping gene *PUM1* in MCF-7 control cells and clones with deleted IRX3 enhancer (Δenh) was measured by qPCR. \**padj* < 0.05, \*\*\**padj* < 0.001; One-way Anova with Holm-Sidak correction for multiple testing. Data are derived from n = 4 biological replicates and representative of two independent experiments. Bar graphs show means ± SD. **C.** Western blot showing the protein levels of IRX3 and protein loading control H3 in MCF-7 control and Δenhancer cells. \**padj* < 0.05, *** *padj* < 0.001; One-Way Anova with Holm-Sidak correction for multiple testing. Bar graphs show means ± SD. **D.** Schematic illustration depicting lentiviral-delivered short hairpin RNA targeting IRX3 transcripts to obtain IRX3 knockdown cells. The figure is created using BioRender. **E.** E. The levels of IRX3 mRNA and protein in MCF-7 control and IRX3 knock-down (shKD) cells were measured by qPCR (left) and western blotting (right), respectively. ***padj < 0.001; One-way Anova with Holm-Sidak correction for multiple testing. Data are derived from n = 3 biological replicates from a single experiment. Bar graphs show means ± SD. **F.** The effect of IRX3 knockdown by either genomic deletion of the upstream enhancer or directly targeting shRNA on global gene expression was assessed by RNA-seq (n=4 biological replicates from a single RNA-seq run). Panther GO – Slim gene ontology analyses (FDR <0.05) was performed on overlapped DEGs (padj < 0.01) in Δenhancer clones and IRX3 knock-down cells (**Supplementary Dataset 1**).

### IRX3 depletion perturbs multiple pathways involved in proliferation, cell survival and tumor suppression

Having established that ERα controls *IRX3* expression via a distal enhancer, we next sought to investigate the functional impact of loss of *IRX3* expression under the control of the characterized enhancer. We first performed RNA sequencing, to assess changes in the global transcriptome of the Δenhancer MCF-7 cell lines and identified 10,475 (5,274 up; 5,201 down) in ΔEnh1 and 10,379 differentially expressed genes (5,096 up; 5,283 down) in ΔEnh3 (*p_adj_* < 0.01) (**Supplementary Dataset 1**). Gene ontology (GO) analyses of the differentially expressed genes between Δenhancer and control cells revealed enrichment of genes involved multiple cancer-related pathways, including EGFR signaling, Ras pathway, p53 signaling, JAK/STAT pathway, cell cycle progression, apoptotic signaling, FAS pathway and angiogenesis (**Supplementary Fig. S5A-D**).

Next, we analyzed MCF-7 cells with lentiviral-delivered shRNAs to assess functional consequences of directly depleting endogenous *IRX3* mRNA (**Figure 3D**). Using two different shRNAs for validation (shKD1 and shKD2), we obtained 70-80% reduction in both IRX3 mRNA and protein levels (**Figure 3E**). RNA sequencing showed 4,726 DEGs (2,528 up; 2,198 down) in shKD1 and 5771 DEGs (2,993 up; 2778 down) in KD2 (*p_adj_* < 0.01) (**Supplementary Dataset 1**). Gene ontology analyses revealed consistent impact of the shRNAs as for Δenhancer cells on many pathways and processes, including EGFR, FAS, integrin, p53, CCKR, cell cycle progression and angiogenesis (**Figure 4F, Supplementary Fig. S6**), with 1,904 differentially expressed genes overlapping between Δenh1, Δenh3, shKD1 and shKD2 (**Supplementary Dataset 1**). Overall, the gene expression data suggests a profound role for IRX3 in MCF-7 proliferation and cell cycle regulation.

**Figure 4.**
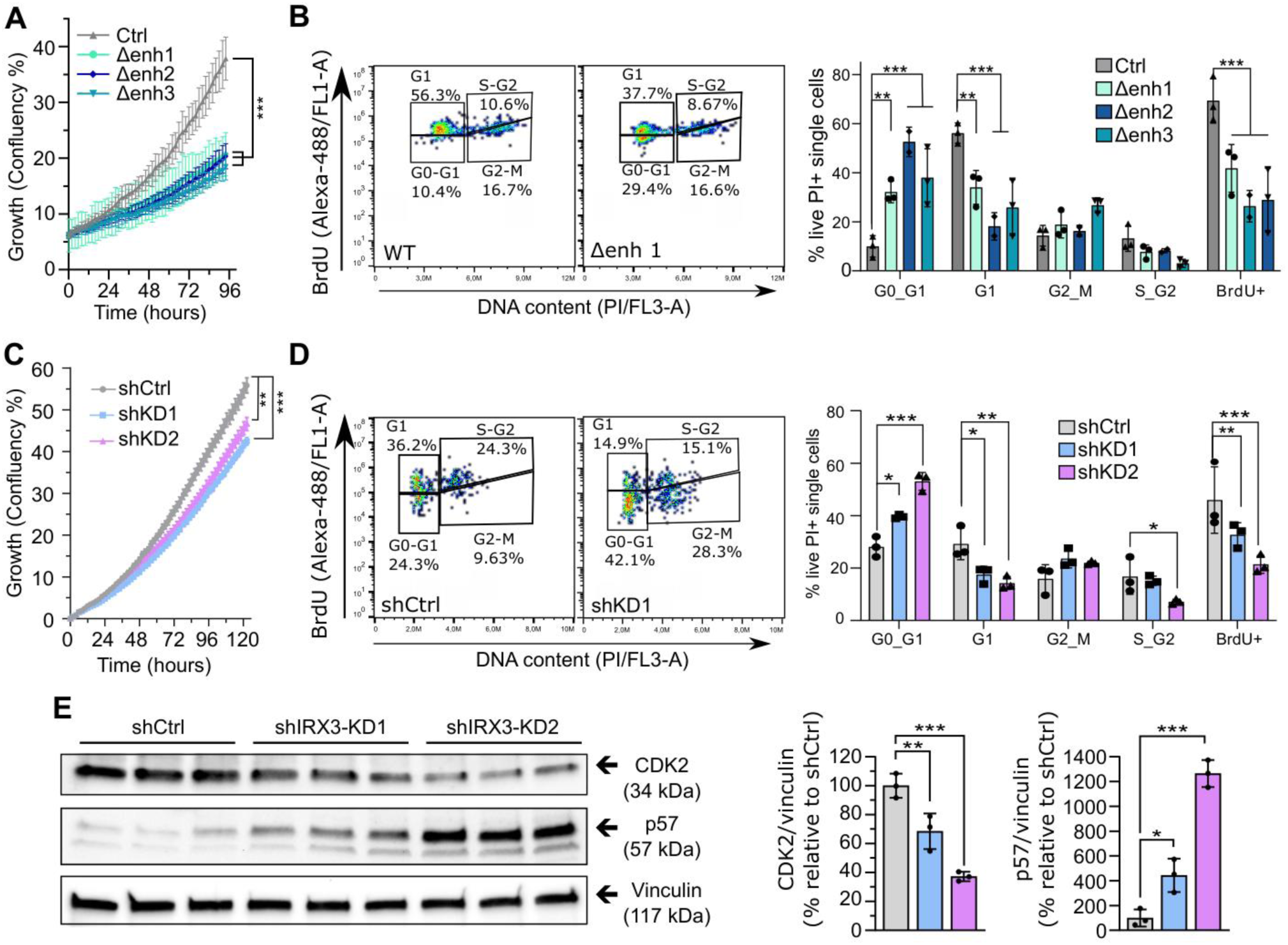
Decreased IRX3 levels reduce cell proliferation through increased G0/G1 cell cycle arrest. **A.** Growth curves of MCF-7 Δenhancer compared to control cells. Cell growth was determined by high content imaging using the IncuCyte S3 Live-Cell system and is presented as percentage of confluence normalized to *t* = 0. Data are derived from n = 8 biological replicates and representative of three independent experiments. \*\**padj* < 0.01, \*\*\**padj* < 0.001; One-Way Anova with Holm-Sidak correction for multiple testing. **B.** Two-dimensional (2D) cell cycle assay, where asynchronous control and Δenhancer MCF-7 cells were pulse-labeled with BrdU for 1 h and left to grow overnight in BrdU-free medium. Harvested cells were fixed and stained with Alexa-488 conjugated anti-BrdU and propidium iodide (PI). Bivariate contour plots of PI vs BrdU from representative samples are shown. The gates represent BrdU-negative cells in G0-G1 and G2-M, and BrdU-positive G1 and S-G2 (left panel). See **Supplementary Fig. S7** for the complete gating strategy. The percentage of cells in each gate is shown in the bar plot (right panel). Data is derived from n = 3 biological replicates from a single experiment. \*\**p_adj_* < 0.01, \*\*\**p_adj_* < 0.001; One-way Anova with Holm-Sidak correction for multiple testing. Bar graphs show means ± SD **C.** Growth curves of MCF-7 breast cancer cells transfected with lentiviral-delivered shRNAs targeting *IRX3*. Cell growth was determined as described in (A). **D.** 2D cell cycle analysis of MCF-7 cells with shRNA-mediated knockdown of *IRX3*. Cells were stained as in (B). Data are derived from n = 3 replicates and representative of three independent experiments. Bivariate contour plots of PI vs BrdU) are shown. The percentage of cells in each gate is shown in the bar plot (right panel). Bar graphs show means ± SD. \*\**padj* < 0.01, \*\*\**padj* < 0.001; One-way Anova with Holm-Sidak correction for multiple testing. **E.** Western blot showing the protein levels of TP57 and CDK2 in MCF-7 control and shKD cells (left panel). Quantification of band intensities, normalized to protein loading control Vinculin (right panel) \**padj* < 0.05, \*\**padj* < 0.01, \*\*\**padj* < 0.001; One-way Anova with Holm-Sidak correction for multiple testing. Bar graphs show means ± SD.

### Reduced IRX3 levels diminish cell proliferation by promoting G0/G1 cell cycle arrest

To functionally assess the effect of IRX3 repression on MCF-7 proliferation, we seeded equal numbers of control, Δenhancer and shKD cells into 96-well plates and monitored their proliferative capacity over time (**Figure 4**). Strikingly, after 4 days, the growth of the Δenhancer cells was reduced by nearly 50% compared to controls (**Figure 4A**). Since GO terms associated with cell cycle regulation were among the top enriched pathways among the DEGs (**Figures 3F, Supplementary Fig. S5, S6, S7A-B, Supplementary Dataset 1**), we additionally performed functional 2D cell cycle analyses using flow cytometry (**Figure 4B**). Here, asynchronous cells were pulse-labelled with BrdU, followed by staining with anti-BrdU and Propidium Iodide (PI) to label cells in the different phases of the cell cycle. We observed a 3 to 5-fold increase in cells arrested in G0/G1 and ∼40-60% reduction in overall BrdU staining in the Δenhancer cells compared to the controls (**Figure 4B**). These findings are in line with our previous observations in preadipocytes (17).

Both the reduced growth rate and increased cell cycle arrest in MCF-7 cells lacking the *IRX3* enhancer were replicated in MCF-7 cells with shKD-mediated repression of *IRX3* compared to controls (**Figure 4C, D**). We further verified these effects in another ER+ cell line, T47D (**Supplementary Fig. S7D-F**). Here, we achieved 80% knockdown of *IRX3* using shRNA (**Supplementary Fig. S7D**), 60% reduction in proliferation (**Supplementary Fig. S7E**), a 2.5-fold increase in G0/G1 arrest and 65% reduction in BrdU staining (**Supplementary Fig. S7F**). Consistent with the observed increase in G0/G1 arrest, immunoblot analyses showed elevated p57 protein levels, accompanied by a corresponding reduction in Cyclin-dependent kinase 2 (CDK2) in IRX3-KD MCF-7 cells relative to controls (**Figure 4E**). Finally, we observed comparable effects on cell cycle progression and proliferation in MCF-7 cells treated with 4OH-tamoxifen or fulvestrant (**Supplementary Fig. S7G**), consistent with previous published findings (42,43). Together, our findings indicate that IRX3 regulates cell cycle progression and proliferation in ER+ breast cancer cells *in vitro*.

### Orthotopic implantation of IRX3-depleted MCF-7 cells in mice increases primary tumor growth and promotes metastatic dissemination

Having established a role for IRX3 in promoting proliferation of ER+ human breast cancer cells *in vitro,* we hypothesized that IRX3 suppression would also abrogate orthotopic tumor growth *in vivo*. To this end, we implanted control and Δenhancer cells, stably expressing both red-shifted firefly luciferase (Red-FLuc) and green fluorescent protein (GFP), into the mammary fat pad of immunocompromised NOD-SCID mice and monitored tumor growth for 7-8 weeks using bioluminescence imaging (**Figure 5A**). Surprisingly, lack of the enhancer and subsequent depletion of IRX3 increased tumor growth, as shown by increased bioluminescent intensity (BLI) and/or tumor weight in IRX3 depleted-recipient mice (**Figure 5B-D**, **Supplementary Figure S8A**). A potent significant increase in metastatic lesions was also found for both Δenhancer clones in the liver, but only significant elevation for Δenh1 in lung and femur compared to control cells (**Figure 5E**).

**Figure 5.**
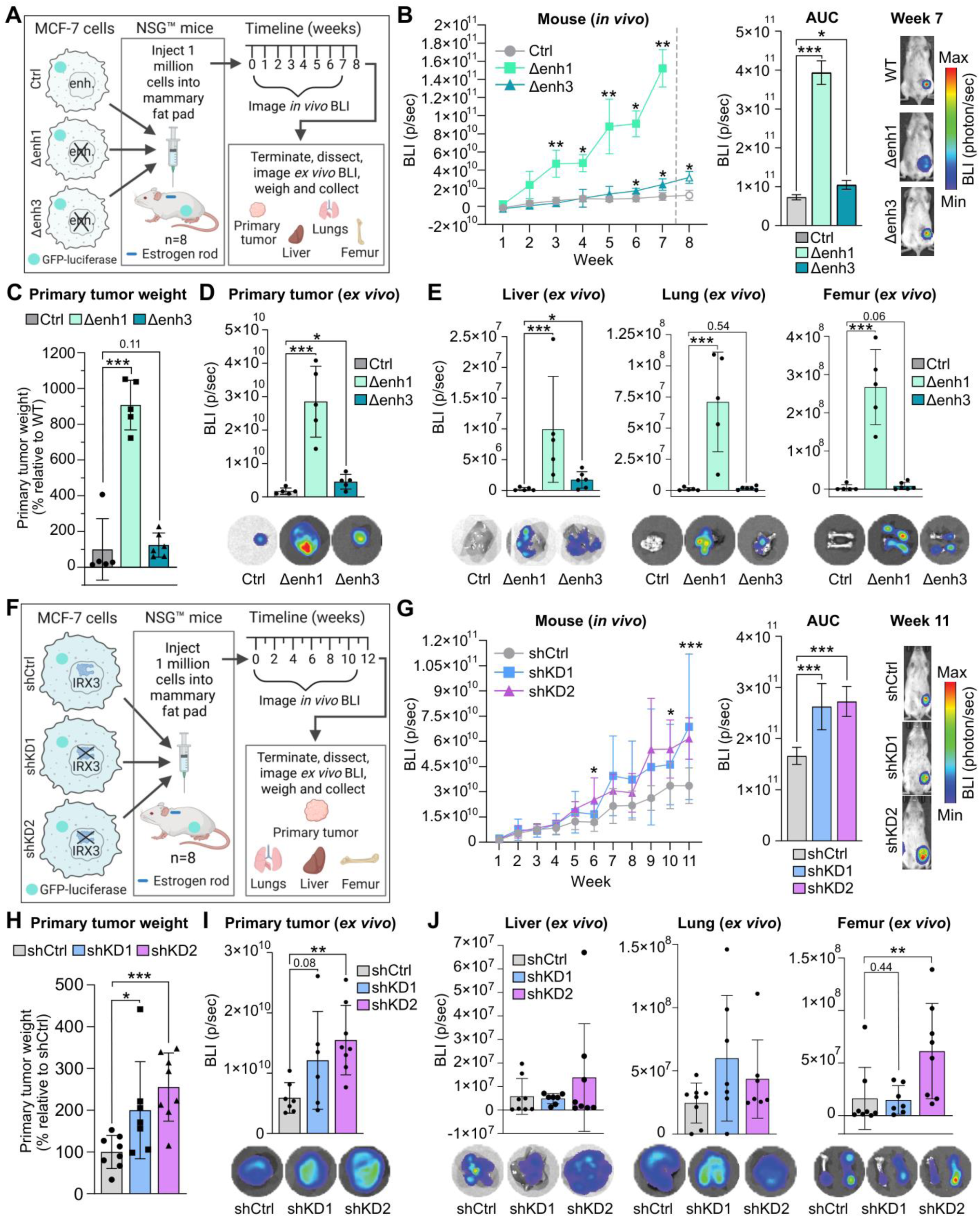
IRX3 depletion in ER+ breast cancer cells increases tumor growth and metastasis *in vivo*. **A.** Schematic illustration of the *in vivo* experiment with MCF-7 cells lacking the *IRX3* enhancer (Δenhancer cells). Control and Δenhancer MCF-7 cells stably expressing a luciferase reporter were orthotopically implanted into the mammary pad of immunocompromised NSG mice (n=8). A rod that supplies human E2 was implanted into the neck region of each mouse 48 h prior to receiving the MCF-7 cells to support the growth of the ER^+^ MCF-7 cells. Tumor growth was monitored by *in vivo* luciferase/bioluminescence imaging (BLI) every week for 7-8 weeks. Mice were then euthanized, and primary tumors, as well as liver, lungs and femur, were imaged *ex vivo* before collection for gene expression and histological analyses. The figure is created in BioRender. **B.** *In vivo* bioluminescence intensity of mice with implanted control and Δenhancer cells on week 1-8 post implantation, n = 5-6 mice from a single experiment (left panel). Data shown as means ± SD. \**p_adj_* < 0.05, \*\**p_adj_* < 0.01; Mixed-effects analysis with Dunnett’s multiple comparisons test was done. Mice implanted with Δenh1 cells were euthanized on week 7 due to very large tumors. Area under the curve (AUC) for each group from week 1-7 (middle panel), data shown as means ± SD. \**p_adj_* < 0.05, \*\*\**p_adj_* < 0.001; Ordinary one-way Anova with Dunnett’s multiple comparisons test was performed. Image of one representative mouse from each group showing bioluminescence intensity of the implanted tumors at week 7 (right panel). **C.** Weight of the excised primary tumors from (B), n = 5-6. Data shown as ± SD. \*\*\**p* < 0.001; One-way Anova with Dunnett’s multiple comparisons was performed. **D.** *Ex vivo* bioluminescence intensity of the excised primary tumors from (B), n = 5. Data shown as means ± SD. \**p_adj_* < 0.05, \*\*\**p_adj_* < 0.001; Ordinary one-way Anova with Dunnett’s multiple comparisons test was done. Representative image of an excised primary tumor from each group showing luciferase intensity (bottom panel). **E.** *Ex vivo* bioluminescence intensity of the metastatic lesions in liver, lung and femurs excised from the mice from (B), n = 5-8. Data shown as means ± SD. \**p_adj_* < 0.05, \*\*\**p_adj_* < 0.001; Ordinary one-way Anova with Dunnett’s multiple comparison test was performed. Representative images of bioluminescence in metastatic lesions in liver, lungs and femurs from each group are shown in the bottom panel. **F.** Schematic illustration of the *in vivo* experiment involving MCF-7 shIRX3 knockdown cells. Control and shIRX3 cells stably expressing a luciferase reporter were implanted into the mammary pad of NSC mice as described in (A), n = 8. The tumors were monitored by bioluminescence imaging for 11 weeks before termination and collection of primary tumors and liver, lungs and femur. The figure is created in BioRender. **G.** *In vivo* bioluminescence intensity of mice with implanted control and shIRX3 cells on week 1-11 post implantation, n = 7-8 mice from a single experiment (left panel). Data shown as means ± SD. \**p_adj_* < 0.05, \*\*\**p_adj_* < 0.001; Mixed-effects analysis with Dunnett’s multiple comparisons test was performed. AUC for each group from week 1-7 (middle panel), data shown as means ± SD. \*\*\**p_adj_* < 0.001; Ordinary one-way Anova with Dunnett’s multiple comparisons test was used. Image of one representative mouse from each group showing bioluminescence intensity of the implanted tumors at week 11 (right panel). **H.** Weight of the excised primary tumors from (G), n = 7-8. Data shown as means ± SD. \**p_adj_* < 0.05, \*\*\**p_adj_* < 0.001; Ordinary one-way Anova with Dunnett’s multiple comparison test was performed. **I.** *Ex vivo* bioluminescence intensity of the excised primary tumors from (G), n = 6-8. Data shown as means ± SD. \*\**p_adj_* < 0.01; Ordinary one-way Anova with Dunnett’s multiple comparison test was used. Representative image of an excised primary tumor from each group showing luciferase intensity (bottom panel). **J.** *Ex vivo* bioluminescence intensity of the metastatic lesions in liver, lung and femurs excised from the mice from (G), n = 7-8. Data shown as means ± SD. \*\**p_adj_* < 0.01; Ordinary one-way Anova Dunnett’s multiple comparison test was performed. Representative images of bioluminescence in metastatic lesions in liver, lungs and femurs from each group are shown in the bottom panel. Data shown as means ± SD.

To confirm that the *in vivo* effects observed in mice implanted with the Δenhancer cells were due to the reduced IRX3 expression *per se*, we performed similar mice experiments with IRX3-KD cells (**Figure 5F**). As for the Δenhancer cells, implantation of the IRX3-KD cells significantly increased tumor growth (**Figure 5G-I**, **Supplementary Figure S8B**). We observed a non-significant increase in metastatic lesions in the liver and femur, whereas lung metastasis was significantly elevated in the shKD1 group only (**Figure 5J**). Histological evaluation and immunohistochemical staining of FFPE tumor sections revealed no changes in mitotic activity or cell proliferation; the latter assessed by Ki-67 staining (**Supplementary Fig. S8C-E**). In contrast, IRX3-suppressed tumors displayed a trend towards increased necrotic areas compared to the controls, based on evaluation of HE staining, respectively (**Supplementary Fig. S8C, F**). Overall, orthotopic xenografts generated from IRX3⍰suppressed cells, either by enhancer knockout or direct shRNA-mediated depletion in MCF-7 cells, consistently exhibited increased tumor growth and enhanced cancer aggressiveness.

### Higher *IRX3* expression in ER+ luminal A and luminal B breast cancer subtypes correlate with improved patient survival

Since repression of *IRX3* was found to have an adverse outcome on ER+ tumor growth in the orthotopic implant model, we next investigated human clinical data for correlations between *IRX3* expression in breast cancer and patient survival. We used the relapse-free survival data in the KMplotter tool and stratified the patients in each subtype to two groups based on median tumor *IRX3* expression (**Figure 6A-B**). Consistent with the orthotopic animal model, lower expression of *IRX3* was associated with reduced relapse-free survival of the patients with ER+ (luminal A and B) tumors (**Figure 6A**). In contrast, no associations were found in HER2+ and basal breast tumor subtypes (**Figure 6A**).

**Figure 6.**
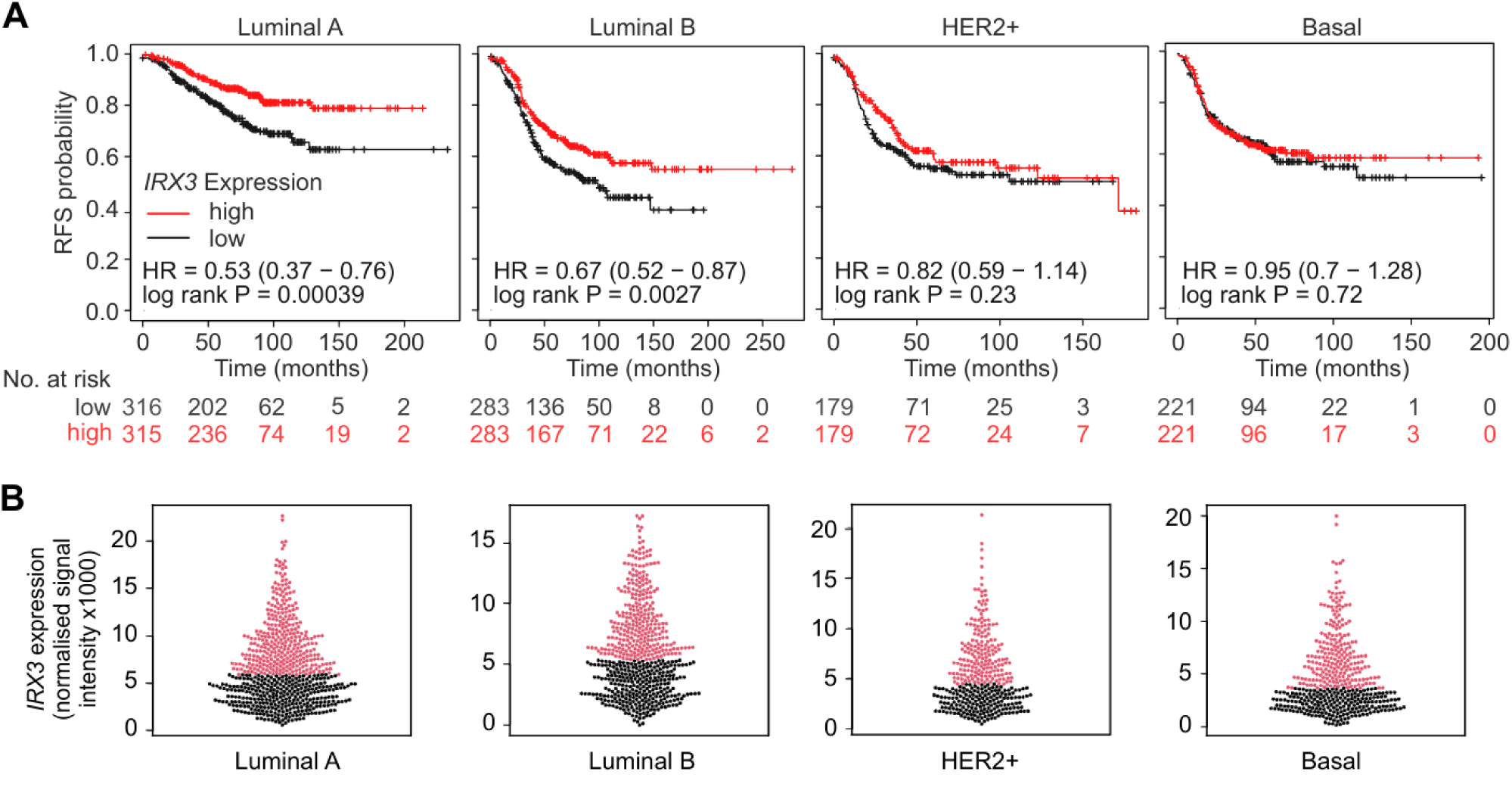
High IRX3 expression is associated with the relapse-free survival in ER+ breast cancer patients. **A.** Association of IRX3 expression with patient survival across the Luminal A, Luminal B, HER2+ and basal breast cancer subtypes. Kaplan-Meier curves for relapse-free survival (RFS) were generated using the KMplotter tool (31) (n = 2032 breast cancer patients). Patients were stratified into high- and low-*IRX3* expression groups based on the median expression level (shown in red and black, respectively). Hazard ratios (HRs) and log-rank test p-values are indicated for each curve. The number of patients at risk at each time point is indicated below each curve. **B.** *IRX3* expression in the breast tumors from the patients shown in A. The data were obtained from the KMplotter tool, which aggregates microarray-based gene expression profiles from multiple studies of patient-derived breast tumors. *IRX3* gene expression from tumors were stratified into high- and low-*IRX3* expression groups based on the median expression level, shown in red and black, respectively.

### Angiogenic programs are activated following IRX3 suppression in ER+ breast tumors

Tumor necrosis is an important hallmark of aggressive cancer types and is associated with increased hypoxia, inflammation, and angiogenesis(30). To further investigate the underlying mechanism in which IRX3 suppression leads to increased tumor growth *in vivo*, we performed RNA sequencing of the excised primary tumors with the marked phenotype of tumor growth (tumors derived from Δenh1 and shKD2 xenografts mice) (**Supplementary Dataset 2**). Further, we found 1023 overlapping up- and down-regulated genes in both Δenh1 and shKD2 primary tumors compared to their respective controls (**Supplementary Dataset 2**). Similar to results from GO analyses on the transcriptomics profile of the IRX3 depleted cells *in vitro* (**Supplementary Fig. S5-6**), GO analysis on the overlapping differentially expressed genes in the primary tumors revealed significant angiogenesis-related terms *in vivo* (**Figure 7A**). Consistent with these findings, gene expression analysis also revealed an elevated VEGF-associated signature, previously reported to reflect enhanced angiogenetic activity in tumors (30) (**Figure 7B, Supplementary Fig. S9A**). To functionally evaluate the angiogenic potential of the IRX3 depleted cells, we utilized a chorioallantoic membrane (CAM) assay *ex ovo.* To this end, shCtrl and shKD2 MCF-7 cells mixed with Matrigel were placed within a Teflon ring on CAM of chicken embryos at day 7 post-fertilization (**Figure 7C, Supplementary Fig. 9B**). After 5 days, the CAMs were dissected and imaged to quantify blood vessels and assess tumor sizes (**Figure 7C-E, Supplementary Fig. S9C-E**). Compared to controls, IRX3 shKD2 MCF-7 cells showed a trend toward forming larger spheroids (region of interest 2) and having enhanced spheroid microcapillary structures and vasculogenic capacity, as indicated by increased vessel area and length (**Figure 7E and Supplementary Fig. S9C**). In contrast, in the negative control area surrounding the spheroid (region of interest 1), there were no trends towards changes in vascularization between shKD2 and shCtrl (**Supplementary Fig. S9C-E**). To further assess whether IRX3 suppression influences angiogenic signaling, we cultured human umbilical vein endothelial cells (HUVECs) in conditioned medium from shIRX3-KD cells and observed a significant increase in HUVEC proliferation *in vitro* (**Figure 7F-G**). Overall, these findings demonstrate that IRX3 plays a role in regulating angiogenetic activity, and consequently, tumor growth in ER+ breast cancer.

**Figure 7.**
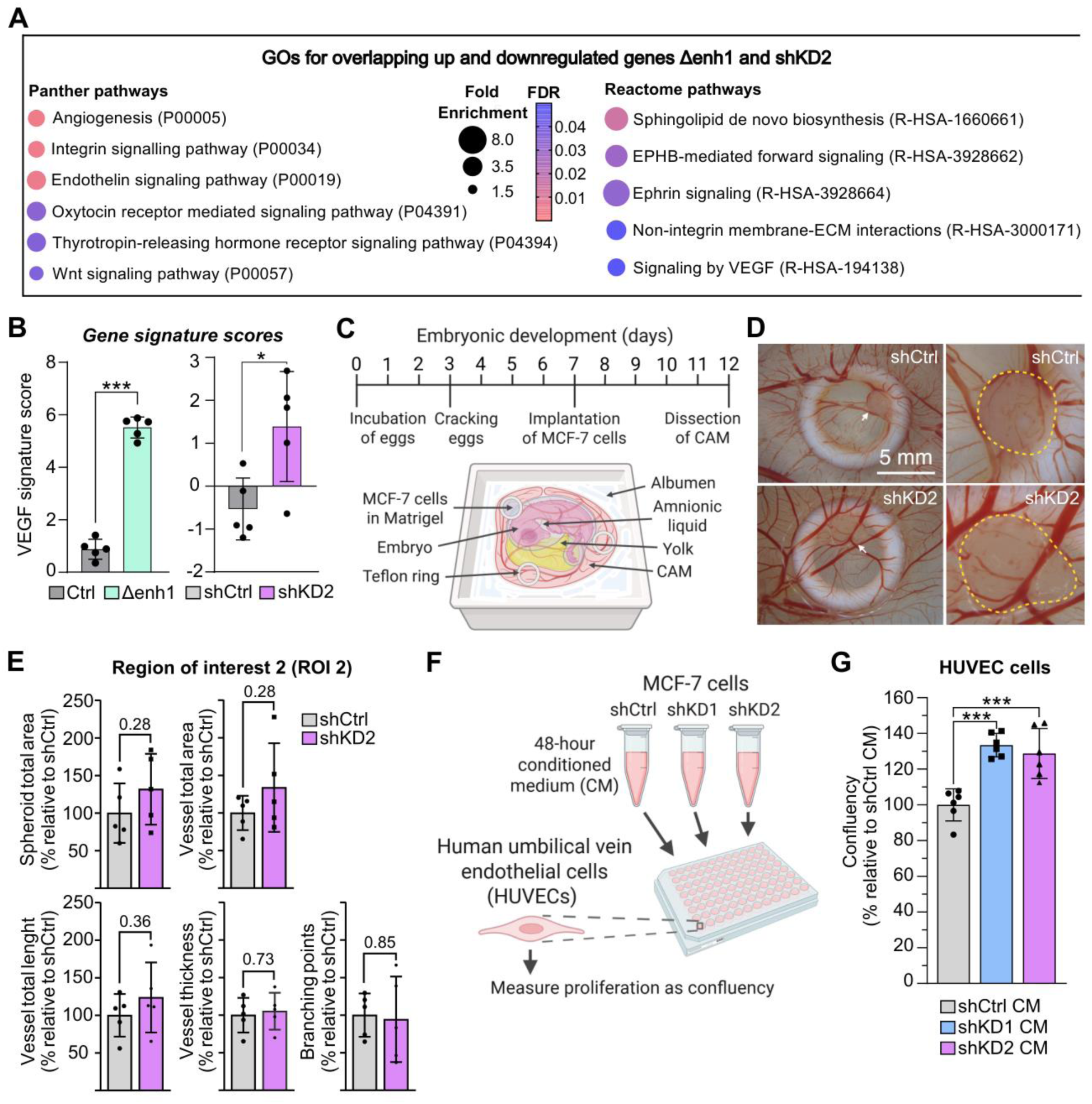
Repression of IRX3 promotes vascularization. **A.** RNA-sequencing of mouse primary tumors (Fig. 5) was performed with n = 5 biological replicates from a single experiment. Overlapping DEGs (*p_adj_* < 0.01) between Δenhancer and control, and shKD2 and shCtrl, were subjected to Panther GO analysis and all significant Panther and Reactome pathways (FDR < 0.05) are shown (**Supplementary Dataset 2**). **B.** VEGF gene signature score in controls, Δenhancer and *IRX3-*KD MCF-7 cells based on the RNA-seq data in (A). The gene signature is calculated using the z-score of the expression of genes within VEGF signature (see Figure S8A) **C.** Schematic illustration of the *ex ovo* chorioallantoic membrane (CAM) assay. At embryonic day 7, MCF-7 shCtrl and *IRX3-*KD cells were prepared in Matrigel and implanted in Teflon rings on top of the CAM of chicken embryos *ex ovo*. After 5 days, the Teflon rings with the CAMs were dissected and imaged prior to quantification of blood vessels by stereomicroscopy and deep-learning imaging analysis. The figure is created in BioRender. **D.** Representative images of dissected CAMs with spheroid-containing Teflon rings. Dashed circles denote the tumor spheroids (region of interest 2 (ROI 2)). Scale bar = 5 mm. **E.** Quantification of blood vessels within region of interest (ROI) 2. Data are derived from n = 5 independent implants in a single experiment. Student’s two-sided t-tests. **F.** Schematic illustration showing conditioned medium from MCF-7 control and *IRX3-*KD cells grown for 48 h added to endothelial HUVEC cells before quantification of HUVEC proliferation. The figure is created in BioRender. **G.** Quantification of HUVEC confluency 4 days after receiving conditioned MCF-7 medium. Data are derived from n = 6 biological replicates from two independent experiments. *** *p_adj_* < 0.001; One-way Anova with Holm-Sidak correction for multiple testing. Data shown as means ± SD.

## Discussion

Given the high prevalence of ER+ breast cancer, the frequent emergence of resistance to endocrine therapies and the poor prognosis upon tumor recurrence, identifying novel molecular drivers that can be therapeutically targeted remains a critical clinical priority. In the present study, we hypothesized an important role for the Iroquois homeobox developmental transcription factor IRX3 in promoting tumor growth and development in ER+ breast cancer. While our *in vitro* data support this hypothesis; IRX3 depletion suppresses proliferation *in vitro;* our *in vivo* data show an opposite role, where orthotopic tumor xenografts with reduced IRX3 levels demonstrated increased angiogenesis and tumor growth compared to controls. These results highlight IRX3 as a context-dependent regulator, whose net effect emerges from a reciprocal interaction between epithelial gene programs and the tumor microenvironment. In support of our findings, a growing literature shows that IRX3 plays diverse, context-dependent roles across cancers. In hematologic malignancies, Somerville et al. (18) and Rahman et al. (44) demonstrated that IRX3 is aberrantly upregulated in acute myeloid leukemia, T-cell acute lymphoblastic leukemia, and B-cell acute lymphoblastic leukemia, where it blocks differentiation and promotes leukemic transformation, together establishing IRX3 as an oncogenic driver in hematopoietic cancers. In contrast, in solid tumors such as gastric (45), colorectal cancer (46), glioblastoma (47), hepatocellular carcinoma (48) and Wilms renal tumors (49), IRX3 is frequently downregulated and interacts with key oncogenic pathways including WNT, Hedgehog, MAPK, TGFβ, PI3K/Akt, and NF-κB, suggesting a tumor-suppressive role. Beyond cancer, previous studies have demonstrated that IRX3 regulates angiogenesis in both developmental and adult contexts (19). Angiogenesis could be driven by recruitment of endothelial progenitor cells and/or by sprouting of pre-existing vasculature (16). Scarlett et al. (50) showed that VEGF-induced IRX3 enhances endothelial migration, Dll4 expression, and tubulogenesis, identifying IRX3 as a pro-angiogenic transcription factor that shapes the vascular microenvironment. Together, these studies establish that IRX3 can function as an oncogene, tumor suppressor, and/or angiogenic modulator depending on tissue context.

In our *in vivo* model, the increased tumor mass was not explained by heightened proliferation, as Ki-67 staining and histopathological assessment did not indicate elevated mitotic activity. Instead, our data indicates that loss of IRX3 triggers compensatory angiogenesis, allowing tumors to overcome reduced epithelial proliferation and ultimately grow larger. Consistent with this, angiogenesis and endothelial–epithelial signaling are known to represent major escape routes through which tumors circumvent the growth-suppressive effects of endocrine therapy, enabling survival in hypoxic niches, enhancing metastatic spread, and reducing drug penetration (51). Similarly, because necrosis – triggered by hypoxia, nutrient deprivation or metabolic stress – can drive angiogenesis, endothelial proliferation, tumor progression and metastasis (52), the increased necrotic regions we observed in IRX3-depleted tumors may have promoted the vascular response. Therefore, the *in vivo* impact of IRX3 loss appears to be driven by an integrated microenvironmental response, encompassing epithelial, endothelial and stromal cues, rather than intrinsic tumor cell behavior alone. This is further consistent with the growing evidence showing that endocrine therapy resistance in ER+ breast cancer arises not only through cell-intrinsic alterations in ER signaling, including ESR1 mutations and downstream pathway rewiring, but also through microenvironment-driven adaptations that enable tumors to sustain growth despite ER blockade (53–55).

In our study, we demonstrate that *IRX3* expression is significantly elevated in ER+ tumors relative to both normal breast tissue and ER- subtypes, and that anti-estrogen treatment in ER+ MCF-7 cells led to reduced *IRX3* mRNA levels. Within breast tumors, we identified a strong co-expression of *IRX3* and *ESR1*, suggesting a tumor-specific association between IRX3 and ER signaling. The association between higher *IRX3* expression and improved relapse-free survival in luminal A and B tumors is consistent with the observation that high ER expression correlates with better overall survival in breast cancer (50), and may, at least in part, reflect increased therapy-responsiveness of tumors with high IRX3 expression, suggesting that IRX3 may also serve as a predictive biomarker for treatment response.

Mechanistically, we identified a previously uncharacterized ERα-bound enhancer, upstream of the *IRX3* locus, that acts as the primary regulatory element controlling *IRX3* expression. Enhancer deletion dramatically suppressed *IRX3,* demonstrating the importance of this regulatory element for *IRX3* expression in ER+ breast cancer cells. Other studies have found the paralog gene *IRX5* to be co-regulated with *IRX3* in some contexts, such as adipose tissue and leukemia (37,38,56). While we found strong chromatin interactions between the enhancer and *IRX5*, this did not translate into functional regulation of *IRX5* as the expression was largely unaffected by deleting the enhancer, suggesting that *IRX3* is the main target. *IRX3* has previously been shown to be regulated by long-range enhancer elements in intron 1 of *FTO* in preadipocytes and neurons (38,57) and intron 8 of *FTO* in acute myeloid leukemia (56). While we found that regions in these introns interacted with IRX3 in ER+ MCF-7 cells, there was no overlap with ERα occupancy. Thus, our findings point to a novel, previously unappreciated enhancer regulating *IRX3* expression.

Interestingly, genome-wide association studies have also identified an association between the SNP rs11075995, located in the second intron of the *FTO* gene, and altered regulation of estrogen biosynthesis genes in ER-negative breast cancer (58). While this SNP was not associated with changes in IRX3 expression in ER-negative breast cancer, the role of this SNP in regulating estrogen production and subsequently estrogen-ERα mediated IRX3 expression in ER+ breast cancer remains to be explored. Previous research in mice found a strong enhancer-mediated upregulation of *Irx3* specifically in female gonads during a critical period of sex determination (59), regulated by canonical Wnt/β-catenin signaling in the fetal ovary (60). Adding to these mechanisms of sexual dimorphism in *Irx3* transcriptional regulation, our data demonstrates that estrogen-ER signaling can regulate *IRX3* in female adults.

Increasing evidence suggests that IRX3 may function as an epigenetic regulator (61), consistent with the broader roles described for homeobox transcription factors in chromatin remodeling and lineage-specific gene regulation (62–65). The IRX-family of proteins, including IRX3, have been implicated in pathways governing developmental patterning, cellular identity, and transcriptional competence (66), as well as processes such as angiogenesis (19) and neurogenesis (67). Moreover, our previous work in preadipocytes demonstrated that IRX3 represses multiple epigenetic regulators and components of the SUMOylation machinery, thereby influencing chromatin modification pathways that regulate cell-cycle progression and differentiation (61). This aligns with additional evidence showing that SUMO-dependent epigenetic mechanisms are linked to mitotic arrest and tumor suppression in breast cancer cells (68), further indicating that IRX3 may interface with chromatin-regulating systems that influence tumor cell proliferation. Taken together, these findings raise the possibility that IRX3’s tumor-modifying effects in ER+ breast cancer may also involve epigenetic modulation of chromatin accessibility, SUMOylation pathways, or enhancer–promoter interactions, thereby contributing to the overall transcriptional landscape that governs tumor growth, angiogenesis, and therapy response. Future studies should address these issues.

Our findings indicate that IRX3-related pathways could be relevant to therapeutic strategies, however, any such approaches would need to carefully account for microenvironmental-associated escape mechanisms activated upon IRX3 loss. Such strategies could include targeting the ERα-dependent enhancer that regulates IRX3, integrating IRX3 inhibition with anti-angiogenic or vascular-normalizing agents, or combining IRX3-directed approaches with established endocrine regimens or CDK4/6 inhibition to prevent microenvironment-mediated resistance (69).

This study has limitations. First, although relevant cell lines and model systems were used to support our conclusions, the principal mechanistic interrogation of the enhancer–IRX3 regulatory axis was conducted in breast cancer cell lines, which may not fully represent the diversity of ER+ breast cancer. Second, both shRNA knockdown and CRISPR-mediated enhancer deletion strategies carry inherent risks of off-target or clone-specific effects that cannot be entirely excluded. Third, the *in vivo* experiments were performed in estrogen-supplemented immunodeficient mice and the *ex ovo* chorioallantoic membrane assay was carried out with modest sample sizes, which may constrain the generalizability of specific observations.

In conclusion, we have established IRX3 as an ERα target gene in ER+ breast cancer cells/tumors and found IRX3 to play a critical role for the growth of these cells/tumors. While IRX3 depletion suppresses epithelial proliferation *in vitro*, it ultimately triggers compensatory angiogenesis *in vivo* to sustain tumor growth and metastatic spread. These findings underscore that cancer progression is governed by interconnected processes, emphasizing the need to consider both cancer cell-intrinsic and microenvironmental effects when targeting pathways in ER+ breast cancer.

## Acknowledgements

We thank Margit Solsvik and Linn Skartveit at the Hormone Laboratory, and Bendik Nordanger and Erling Høivik at the Department of Pathology, Haukeland University Hospital for their excellent technical contribution and insightful advice. We also acknowledge the Flow Cytometry Core Facility, Department of Clinical Science, the Molecular Imaging Center (MIC), Department of Biomedicine, and the Genomics Core Facility (GCF), University of Bergen. GCF is a part of the NorSeq consortium and is supported by major grants from the Research Council of Norway (grant no. 245979/F50) and the Bergen Research Foundation (BFS) (grant no. BFS2017TMT04 and BFS2017TMT08).

## Funding

This work was funded by grants received from the Western Norway Regional Health Authority, the Trond Mohn Foundation (BFS2017NUTRITIONLAB), University of Bergen and Haukeland University Hospital.

## Authors’ Contributions

Conception and design: P.P.S, JI.B, R.Å.J, S.N.D, G.M.

Development of methodology: P.P.S, JI.B, R.Å.J, S.Y, K.F

Acquisition of data (provided animals, acquired and managed patients, provided facilities, etc.): P.P.S, JI.B, R.Å.J, M.P, S.Y, K.F, E.W

Analysis and interpretation of data (e.g., statistical analysis, biostatistics, computational analysis): P.P.S, JI.B, R.Å.J, M.P, S.Y, K.F, E.W, S.N.D, G.M

Writing, review, and/or revision of the manuscript: P.P.S, JI.B, R.Å.J, M.P, E.M, S.Y, K.M, E.W, S.N.D, G.M

Administrative, technical, or material support (i.e., reporting or organizing data, constructing databases): P.P.S, JI.B, R.Å.J, S.Y, K.F

Study supervision: G.M.

## Notes

The authors declare no potential conflicts of interest.

### Competing Interest Statement

The authors have declared no competing interest.

## References

1. Siegel RL, Kratzer TB, Giaquinto AN, Sung H, Jemal A. Cancer statistics, 2025. CA Cancer J Clin 2025;75:10–45

2. Fishman J, Osborne MP, Telang NT. The role of estrogen in mammary carcinogenesis. Ann N Y Acad Sci 1995;768:91–100

3. Harbeck N, Penault-Llorca F, Cortes J, Gnant M, Houssami N, Poortmans P, et al. Breast cancer. Nat Rev Dis Primers 2019;5:66

4. Yager JD, Davidson NE. Estrogen carcinogenesis in breast cancer. N Engl J Med 2006;354:270–82

5. Lloyd MC, Alfarouk KO, Verduzco D, Bui MM, Gillies RJ, Ibrahim ME, et al. Vascular measurements correlate with estrogen receptor status. BMC Cancer 2014;14:279

6. Mueller MD, Vigne JL, Minchenko A, Lebovic DI, Leitman DC, Taylor RN. Regulation of vascular endothelial growth factor (VEGF) gene transcription by estrogen receptors alpha and beta. Proc Natl Acad Sci U S A 2000;97:10972–7

7. Risau W. Mechanisms of angiogenesis. Nature 1997;386:671–4

8. Suriano R, Chaudhuri D, Johnson RS, Lambers E, Ashok BT, Kishore R, et al. 17Beta-estradiol mobilizes bone marrow-derived endothelial progenitor cells to tumors. Cancer Res 2008;68:6038–42

9. Losordo DW, Isner JM. Estrogen and angiogenesis: A review. Arterioscler Thromb Vasc Biol 2001;21:6–12

10. Arimidex TAoiCTG, Forbes JF, Cuzick J, Buzdar A, Howell A, Tobias JS, et al. Effect of anastrozole and tamoxifen as adjuvant treatment for early-stage breast cancer: 100-month analysis of the ATAC trial. Lancet Oncol 2008;9:45–53

11. Early Breast Cancer Trialists’ Collaborative G, Davies C, Godwin J, Gray R, Clarke M, Cutter D, et al. Relevance of breast cancer hormone receptors and other factors to the efficacy of adjuvant tamoxifen: patient-level meta-analysis of randomised trials. Lancet 2011;378:771–84

12. Pan H, Gray R, Braybrooke J, Davies C, Taylor C, McGale P, et al. 20-Year Risks of Breast-Cancer Recurrence after Stopping Endocrine Therapy at 5 Years. N Engl J Med 2017;377:1836–46

13. Richman J, Dowsett M. Beyond 5 years: enduring risk of recurrence in oestrogen receptor-positive breast cancer. Nat Rev Clin Oncol 2019;16:296–311

14. Ali S, Coombes RC. Endocrine-responsive breast cancer and strategies for combating resistance. Nat Rev Cancer 2002;2:101–12

15. Bianchini G, Pusztai L, Karn T, Iwamoto T, Rody A, Kelly C, et al. Proliferation and estrogen signaling can distinguish patients at risk for early versus late relapse among estrogen receptor positive breast cancers. Breast Cancer Res 2013;15:R86

16. Helland T, Gjerde J, Dankel S, Fenne IS, Skartveit L, Drangevag A, et al. The active tamoxifen metabolite endoxifen (4OHNDtam) strongly down-regulates cytokeratin 6 (CK6) in MCF-7 breast cancer cells. PLoS One 2015;10:e0122339

17. Bjune JI, Dyer L, Rosland GV, Tronstad KJ, Njolstad PR, Sagen JV, et al. The homeobox factor Irx3 maintains adipogenic identity. Metabolism 2020;103:154014

18. Somerville TDD, Simeoni F, Chadwick JA, Williams EL, Spencer GJ, Boros K, et al. Derepression of the Iroquois Homeodomain Transcription Factor Gene IRX3 Confers Differentiation Block in Acute Leukemia. Cell Rep 2018;22:638–52

19. Scarlett K, Pattabiraman V, Barnett P, Liu D, Anderson LM. The proangiogenic effect of iroquois homeobox transcription factor Irx3 in human microvascular endothelial cells. J Biol Chem 2015;290:6303–15

20. Brown RM, Wang L, Fu A, Kannan A, Mussar M, Bagchi IC, et al. Irx3 promotes gap junction communication between uterine stromal cells to regulate vascularization during embryo implantationdagger. Biol Reprod 2022;106:1000–10

21. Gaborit N, Sakuma R, Wylie JN, Kim KH, Zhang SS, Hui CC, et al. Cooperative and antagonistic roles for Irx3 and Irx5 in cardiac morphogenesis and postnatal physiology. Development 2012;139:4007–19

22. Liu H, Naismith JH. An efficient one-step site-directed deletion, insertion, single and multiple-site plasmid mutagenesis protocol. BMC Biotechnol 2008;8:91

23. Anzalone AV, Randolph PB, Davis JR, Sousa AA, Koblan LW, Levy JM, et al. Search-and-replace genome editing without double-strand breaks or donor DNA. Nature 2019;576:149–57

24. Bankhead P, Loughrey MB, Fernandez JA, Dombrowski Y, McArt DG, Dunne PD, et al. QuPath: Open source software for digital pathology image analysis. Sci Rep 2017;7:16878

25. Ritchie ME, Phipson B, Wu D, Hu Y, Law CW, Shi W, et al. limma powers differential expression analyses for RNA-sequencing and microarray studies. Nucleic Acids Res 2015;43:e47

26. Pal B, Chen Y, Vaillant F, Capaldo BD, Joyce R, Song X, et al. A single-cell RNA expression atlas of normal, preneoplastic and tumorigenic states in the human breast. EMBO J 2021;40:e107333

27. Hao Y, Stuart T, Kowalski MH, Choudhary S, Hoffman P, Hartman A, et al. Dictionary learning for integrative, multimodal and scalable single-cell analysis. Nat Biotechnol 2024;42:293–304

28. Love MI, Huber W, Anders S. Moderated estimation of fold change and dispersion for RNA-seq data with DESeq2. Genome Biol 2014;15:550

29. Thomas PD, Ebert D, Muruganujan A, Mushayahama T, Albou LP, Mi H. PANTHER: Making genome-scale phylogenetics accessible to all. Protein Sci 2022;31:8–22

30. Hu Z, Fan C, Livasy C, He X, Oh DS, Ewend MG, et al. A compact VEGF signature associated with distant metastases and poor outcomes. BMC Med 2009;7:9

31. Posta M, Gyorffy B. Pathway-level mutational signatures predict breast cancer outcomes and reveal therapeutic targets. Br J Pharmacol 2025;182:5734–47

32. Curtis C, Shah SP, Chin SF, Turashvili G, Rueda OM, Dunning MJ, et al. The genomic and transcriptomic architecture of 2,000 breast tumours reveals novel subgroups. Nature 2012;486:346–52

33. Hintzsche J, Kim J, Yadav V, Amato C, Robinson SE, Seelenfreund E, et al. IMPACT: a whole-exome sequencing analysis pipeline for integrating molecular profiles with actionable therapeutics in clinical samples. J Am Med Inform Assoc 2016;23:721–30

34. Castro-Mondragon JA, Riudavets-Puig R, Rauluseviciute I, Lemma RB, Turchi L, Blanc-Mathieu R, et al. JASPAR 2022: the 9th release of the open-access database of transcription factor binding profiles. Nucleic Acids Res 2022;50:D165–D73

35. Welboren WJ, van Driel MA, Janssen-Megens EM, van Heeringen SJ, Sweep FC, Span PN, et al. ChIP-Seq of ERalpha and RNA polymerase II defines genes differentially responding to ligands. EMBO J 2009;28:1418–28

36. Ross-Innes CS, Stark R, Teschendorff AE, Holmes KA, Ali HR, Dunning MJ, et al. Differential oestrogen receptor binding is associated with clinical outcome in breast cancer. Nature 2012;481:389–93

37. Smemo S, Tena JJ, Kim KH, Gamazon ER, Sakabe NJ, Gomez-Marin C, et al. Obesity-associated variants within FTO form long-range functional connections with IRX3. Nature 2014;507:371–5

38. Claussnitzer M, Dankel SN, Kim KH, Quon G, Meuleman W, Haugen C, et al. FTO Obesity Variant Circuitry and Adipocyte Browning in Humans. N Engl J Med 2015;373:895–907

39. Ragvin A, Moro E, Fredman D, Navratilova P, Drivenes O, Engstrom PG, et al. Long-range gene regulation links genomic type 2 diabetes and obesity risk regions to HHEX, SOX4, and IRX3. Proc Natl Acad Sci U S A 2010;107:775–80

40. Zwart W, Theodorou V, Kok M, Canisius S, Linn S, Carroll JS. Oestrogen receptor-co-factor-chromatin specificity in the transcriptional regulation of breast cancer. EMBO J 2011;30:4764–76

41. Shang Y, Hu X, DiRenzo J, Lazar MA, Brown M. Cofactor dynamics and sufficiency in estrogen receptor-regulated transcription. Cell 2000;103:843–52

42. Dolfi SC, Jager AV, Medina DJ, Haffty BG, Yang JM, Hirshfield KM. Fulvestrant treatment alters MDM2 protein turnover and sensitivity of human breast carcinoma cells to chemotherapeutic drugs. Cancer Lett 2014;350:52–60

43. Otto AM, Paddenberg R, Schubert S, Mannherz HG. Cell-cycle arrest, micronucleus formation, and cell death in growth inhibition of MCF-7 breast cancer cells by tamoxifen and cisplatin. J Cancer Res Clin Oncol 1996;122:603–12

44. Rahman S, Bloye G, Farah N, Demeulemeester J, Costa JR, O’Connor D, et al. Focal deletions of a promoter tether activate the IRX3 oncogene in T-cell acute lymphoblastic leukemia. Blood 2024;144:2319–26

45. Dos Santos EC, Petrone I, Binato R, Abdelhay E. Iroquois Family Genes in Gastric Carcinogenesis: A Comprehensive Review. Genes (Basel) 2023;14

46. Martorell O, Barriga FM, Merlos-Suarez A, Stephan-Otto Attolini C, Casanova J, Batlle E, et al. Iro/IRX transcription factors negatively regulate Dpp/TGF-beta pathway activity during intestinal tumorigenesis. EMBO Rep 2014;15:1210–8

47. Gao Y, Zhang G, Yu Y, Gao J, Ren S, Wei X, et al. IRX3-CDK14 axis promotes glioblastoma progression by regulating LRP6-mediated canonical Wnt/beta-catenin pathway. Cell Death Dis 2025;17:127

48. Wang P, Zhuang C, Huang D, Xu K. Downregulation of miR-377 contributes to IRX3 deregulation in hepatocellular carcinoma. Oncol Rep 2016;36:247–52

49. Holmquist Mengelbier L, Lindell-Munther S, Yasui H, Jansson C, Esfandyari J, Karlsson J, et al. The Iroquois homeobox proteins IRX3 and IRX5 have distinct roles in Wilms tumour development and human nephrogenesis. J Pathol 2019;247:86–98

50. Barron-Gallardo CA, Garcia-Chagollan M, Moran-Mendoza AJ, Delgadillo-Cristerna R, Martinez-Silva MG, Villasenor-Garcia MM, et al. A gene expression signature in HER2+ breast cancer patients related to neoadjuvant chemotherapy resistance, overall survival, and disease-free survival. Front Genet 2022;13:991706

51. Gacche RN. Compensatory angiogenesis and tumor refractoriness. Oncogenesis 2015;4:e153

52. Karsch-Bluman A, Feiglin A, Arbib E, Stern T, Shoval H, Schwob O, et al. Tissue necrosis and its role in cancer progression. Oncogene 2019;38:1920–35

53. Diaz Bessone MI, Gattas MJ, Laporte T, Tanaka M, Simian M. The Tumor Microenvironment as a Regulator of Endocrine Resistance in Breast Cancer. Front Endocrinol (Lausanne) 2019;10:547

54. Yuan J, Yang L, Li Z, Zhang H, Wang Q, Huang J, et al. The role of the tumor microenvironment in endocrine therapy resistance in hormone receptor-positive breast cancer. Front Endocrinol (Lausanne) 2023;14:1261283

55. Belachew EB, Sewasew DT. Molecular Mechanisms of Endocrine Resistance in Estrogen-Positive Breast Cancer. Front Endocrinol (Lausanne) 2021;12:599586

56. Camera F, Romero-Camarero I, Revell BH, Amaral FMR, Sinclair OJ, Simeoni F, et al. Differentiation block in acute myeloid leukemia regulated by intronic sequences of FTO. iScience 2023;26:107319

57. Sobreira DR, Joslin AC, Zhang Q, Williamson I, Hansen GT, Farris KM, et al. Extensive pleiotropism and allelic heterogeneity mediate metabolic effects of IRX3 and IRX5. Science 2021;372:1085–91

58. Wiggins GAR, Black MA, Dunbier A, Merriman TR, Pearson JF, Walker LC. Variable expression quantitative trait loci analysis of breast cancer risk variants. Sci Rep 2021;11:7192

59. Jorgensen JS, Gao L. Irx3 is differentially up-regulated in female gonads during sex determination. Gene Expr Patterns 2005;5:756–62

60. Koth ML, Garcia-Moreno SA, Novak A, Holthusen KA, Kothandapani A, Jiang K, et al. Canonical Wnt/beta-catenin activity and differential epigenetic marks direct sexually dimorphic regulation of Irx3 and Irx5 in developing mouse gonads. Development 2020;147

61. Bjune JI, Laber S, Lawrence-Archer L, Nothnagel PMC, Yamada S, Zhao X, et al. IRX3 controls a SUMOylation-dependent differentiation switch in adipocyte precursor cells. Nat Commun 2025;16:7248

62. Magli A, Baik J, Mills LJ, Kwak IY, Dillon BS, Mondragon Gonzalez R, et al. Time-dependent Pax3-mediated chromatin remodeling and cooperation with Six4 and Tead2 specify the skeletal myogenic lineage in developing mesoderm. PLoS Biol 2019;17:e3000153

63. Gordon JA, Hassan MQ, Saini S, Montecino M, van Wijnen AJ, Stein GS, et al. Pbx1 represses osteoblastogenesis by blocking Hoxa10-mediated recruitment of chromatin remodeling factors. Mol Cell Biol 2010;30:3531–41

64. Desanlis I, Kherdjemil Y, Mayran A, Bouklouch Y, Gentile C, Sheth R, et al. HOX13-dependent chromatin accessibility underlies the transition towards the digit development program. Nat Commun 2020;11:2491

65. Cain B, Gebelein B. Mechanisms Underlying Hox-Mediated Transcriptional Outcomes. Front Cell Dev Biol 2021;9:787339

66. Goutam RS, Kaushik N, Kumar B, Jung W, Kumar S, Lee SH, et al. Multifaceted role of Iroquois signaling in development and diseases. Mol Cells 2025;48:100296

67. Catela C, Assimacopoulos S, Chen Y, Tsioras K, Feng W, Kratsios P. The Iroquois (Iro/Irx) homeobox genes are conserved Hox targets involved in motor neuron development. iScience 2025;28:112210

68. Eifler K, Cuijpers SAG, Willemstein E, Raaijmakers JA, El Atmioui D, Ovaa H, et al. SUMO targets the APC/C to regulate transition from metaphase to anaphase. Nat Commun 2018;9:1119

69. Hanker AB, Sudhan DR, Arteaga CL. Overcoming Endocrine Resistance in Breast Cancer. Cancer Cell 2020;37:496–513

