## Supplementary Information for "ERα-regulated IRX3 controls the growth of ER-positive breast tumors"

**Supplementary Table**

**Supplementary Table S1. Sequences of primers and sgRNAs used for cloning, mutagenesis, real-time PCR and genomic deletion of the enhancer**

| **Name** | **Sequence** | **Application** |
| --- | --- | --- |
| hIRX3-enh_2kb_BglII_F | 5’*-AAAA*AGATCTACTGTGGGGATTTGGTGTGC-3’ | Cloning |
| hIRX3-enh_1.5kb_BglII_F | 5’-*AAAA*AGATCTGGCCTGGACTCGTGAGC-3’ | Cloning |
| hIRX3-enh_0.8kb_BglII_F | 5’-*AAAA*AGATCTGCCAAGTTGTTTACACTGACCAT-3’ | Cloning |
| hIRX3-enh_all_HindIII_R | 5’-*TAATAAGCTT*TAGCACCGCAGAGAACGC-3’ | Cloning |
| RVprimer 3_F | 5’-CTAGCAAAATAGGCTGTCCC-3’ | Colony PCR |
| hIRX3-enh_ col.PCR_R | 5’-TGGGGAGAGCCGATAAGAGG-3’ | Colony PCR |
| ΔERE1_mut_F | 5’-TTGCAGCATTGTTTGTTAAATAATGAACTAATCGGGTCATCTTATGGGAGAGCTGGGC-3’ | Mutagenesis |
| ΔERE1_mut_R | 5’-GATTAGTTCATTATTTAACAAACAATGCTGCAACTGCTCATGAAAACCACAAAACAGGCTG-3’ | Mutagenesis |
| ΔERE4_Del_F | 5’-GGCCAGGAGGGCATCAGCACCAGC-3’ | Mutagenesis |
| ΔERE4_Del_R | 5’-GCTGGTGCTGATGCCCTCCTGGCC-3’ | Mutagenesis |
| ΔERE1,5_Del_F | 5’-GCAGTTGCAGCATTGTTTGTTTCTTATGGGAGAGCTG-3’ | Mutagenesis |
| ΔERE1,5_Del_R | 5’-CAGCTCTCCCATAAGAAACAAACAATGCTGCAACTGC-3’ | Mutagenesis |
| ΔERE6_del_F | 5’-CCTCTTTTGAAAGCAGCCGTGGAGAAGGAATGGG-3’ | Mutagenesis |
| ΔERE6_del_R | 5’-CCCATTCCTTCTCCACGGCTGCTTTCAAAAGAGG-3’ | Mutagenesis |
| ΔERE7_del_F | 5’-CTGTGCAGGTGGCTCTAATCCTGCTGCTTCTA-3’ | Mutagenesis |
| ΔERE7_del_R | 5’-TAGAAGCAGCAGGATTAGAGCCACCTGCACAG-3’ | Mutagenesis |
| ΔERE8_del_F | 5’-CAATGCAGTTCTAGCCCTCATAAAATGACTATTTTTCCCTGG-3’ | Mutagenesis |
| ERE8_del_R | 5’-CCAGGGAAAAATAGTCATTTTATGAGGGCTAGAACTGCATTG-3’ | Mutagenesis |
| ΔERE9_del_F | 5’-AAGATGAGTCTTTTCTAAATCTTAGTGGGATGAACTCTTACAAAAGTC-3’ | Mutagenesis |
| ΔERE9_del_R | 5’-GACTTTTGTAAGAGTTCATCCCACTAAGATTTAGAAAAGACTCATCTT-3’ | Mutagenesis |
| ΔERE2_del_F | 5’-GACCTCCTCTTGGTGCCAAAGATGGAATGGAGAT-3’ | Mutagenesis |
| ΔERE2_del_R | 5’-ATCTCCATTCCATCTTTGGCACCAAGAGGAGGTC-3’ | Mutagenesis |
| ΔERE10_del_F | 5’-TCTCTTTAAAAGCCAAGTTGTTTACACATATTTACTCTCAACTTCTAATTATTG-3’ | Mutagenesis |
| ΔERE10_del_R | 5’-CAATAATTAGAAGTTGAGAGTAAATATGTGTAAACAACTTGGCTTTTAAAGAGA-3’ | Mutagenesis |
| ΔERE11_del_F | 5’-AGTTTTGCTCCCTAGCTAGAGACATTGCTCCAACC-3’ | Mutagenesis |
| ΔERE11_del_R | 5’-GGTTGGAGCAATGTCTCTAGCTAGGGAGCAAAACT-3’ | Mutagenesis |
| ΔERE3_del_F | 5’-GCTCTGCCCTGCTACTCAGAAGTGAGCTTCCG-3’ | Mutagenesis |
| ΔERE3_del_R | 5’-CGGAAGCTCACTTCTGAGTAGCAGGGCAGAGC-3’ | Mutagenesis |
| 5’ sgRNA (top oligo) | 5’-**CACC**GCTAATGGAGCTAAGTCACG-3’ | Genomic deletion of the enhancer |
| 5’ sgRNA (bottom oligo) | 5’-**AAAC**CGTGACTTAGCTCCATTAGC-3’ | Genomic deletion of the enhancer |
| 3’ sgRNA (top oligo) | 5’-**CACC**GCAGGGCATTGTTCCTAAGG-3’ | Genomic deletion of the enhancer |
| 3’ sgRNA (bottom oligo) | 5’-**AAAC**CCTTAGGAACAATGCCCTGC-3’ | Genomic deletion of the enhancer |
| Outside forward primer | 5’-AGACTGCCTCTCAAGGAATGA-3’ | Enhancer screening primer |
| Outside reverse primer | 5’-GAGTGCTGGGGTGTGTTTG-3’ | Enhancer screening primer |
| Inside forward primer | 5’-TTTTCCTTCTCTCCTTCTGCCC-3’ | Enhancer screening primer |
| Inside reverse primer | 5’-AAATACAGTCAGGTAGGTTTCGT-3’ | Enhancer screening primer |
| Forward flanking primer | 5’-GGGAGATAAGGCGAGGGAGA-3’ | Enhancer screening primer |
| Reverse flanking primer | 5’-TGACCCAACAAACAATGCTG-3’ | Enhancer screening primer |
| IRX3-F | 5’-TCACCAAGATGACCCTCACC-3’ | Real-time PCR |
| IRX3-R | 5’-CCTCCTCTTCGTCTTCCTCC-3’ | Real-time PCR |
| TBP-F | 5’-TGAATCTTGGTTGTAAACTTGACC-3’ | Real-time PCR |
| TBP-R | 5’-CTCATGATTACCGCAGCAAA-3’ | Real-time PCR |
| PUM1-F | 5’-TCACATGGATCCTCTTCAAGC-3’ | Real-time PCR |
| PUM1_R | 5’-CCTGGAGCAGCAGAGATGTAT-3’ | Real-time PCR |

Nucleotides in italics presents sitting sequences; Underlined sequences are BglII RE site; Italics and underlined are HindIII RE site; bold sequences are cloning overhangs for sgRNA oligos. sgRNA, single guide RNA

**Supplementary Figures**

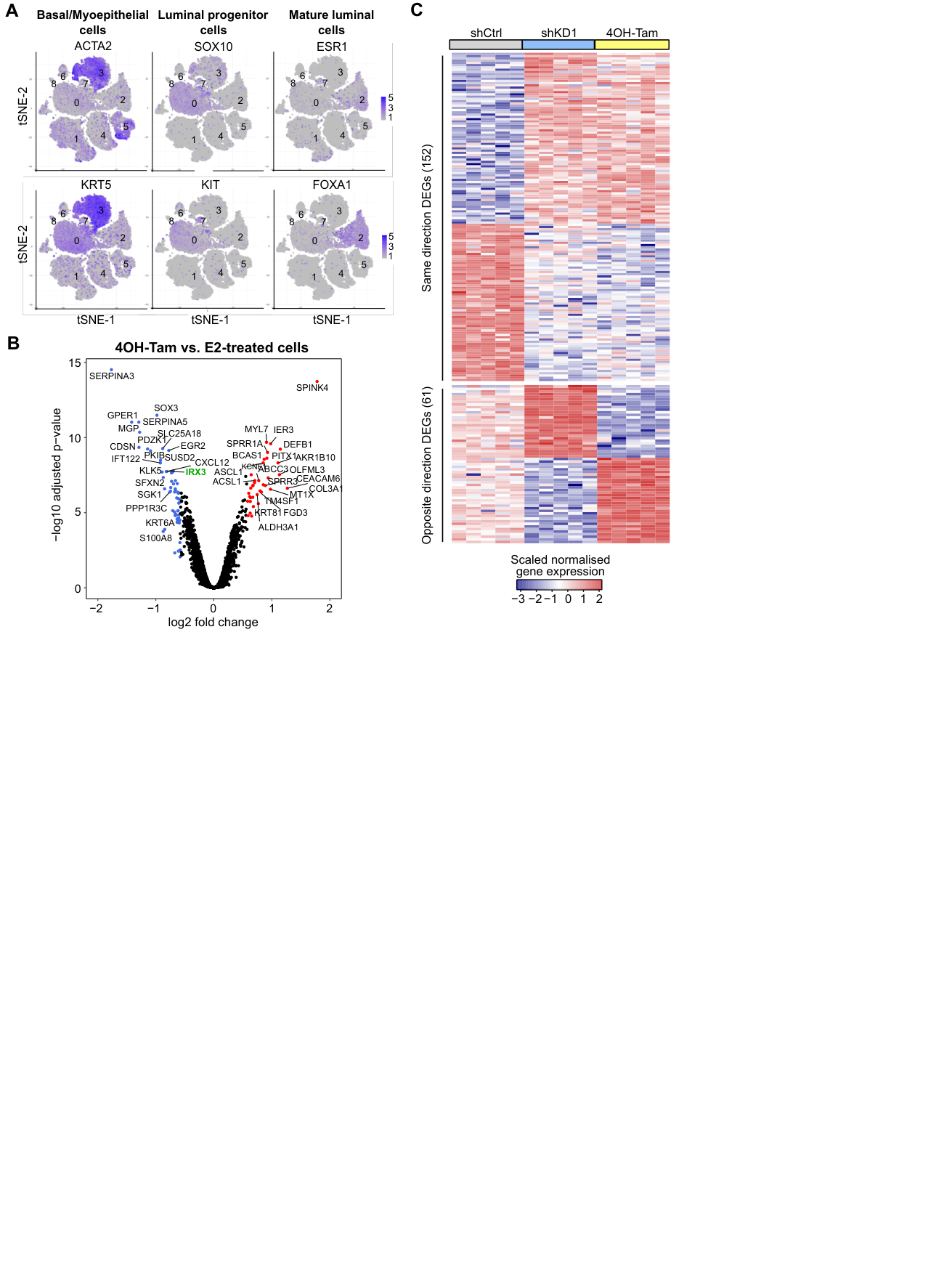

**Figure S1. *IRX3* is highly expressed in epithelial cells and regulates ERα-dependent genes**

A. The expression of specific markers in basal/myoepithelial cells (ACTA2, KRT5, cluster 3), luminal progenitor cells (SOX10 and KIT, cluster 0), and mature luminal cells (ESR1 and FAOXA1, cluster 2) represent distinct epithelial subpopulations within normal breast tissues shown as tSNE plots. The data are derived from publicly available scRNA-seq datasets (1) (GSE161529).
B. Volcano plot of the differentially expressed genes (DEGs) comparing MCF-7 cells that were treated with 4OH-Tam and E2. The data are derived from a previous microarray expression dataset (2) (E-MTAB-2729) with n$=$5 biological replicates.
C. MCF-7 cells were treated with either non-targeting short hairpin control (shCtrl), shRNA targeting *IRX3* (shKD1) or 250 nM 4OH-Tamoxifen (4OH-Tam) (n $=$ 5 biological replicates). Global changes in gene expression were assessed by RNA-sequencing. Heatmap of overlapping differentially expressed genes (DEGs) (n=152, DEGs similar direction and n=61, DEGs opposite direction) is shown.

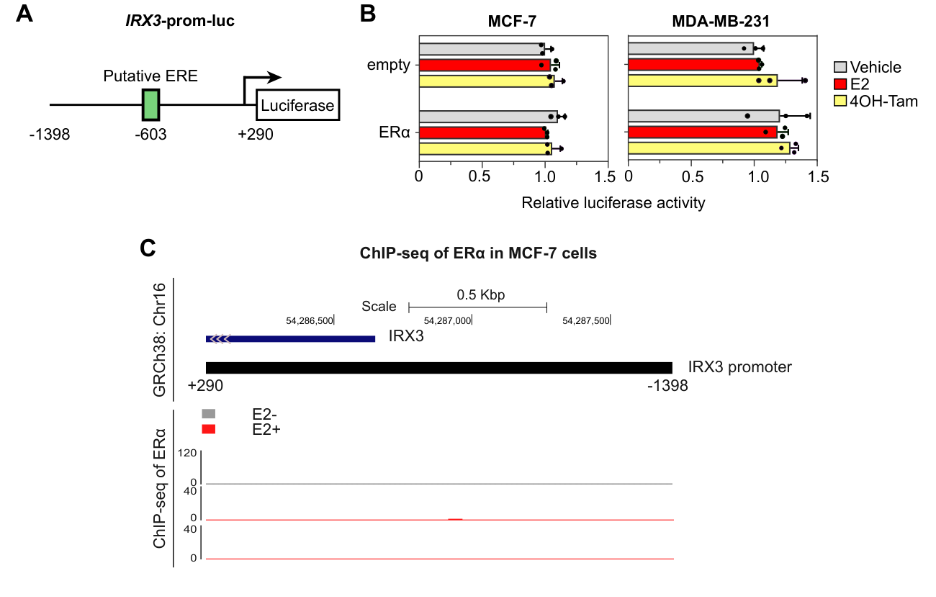
**Figure S2. ERα does not activate the IRX3 promoter *in vitro***

A. Schematic view of the *IRX3* promoter in a luciferase reporter vector (*IRX3*-prom-luc). The position of a JASPAR-predicted ERα response element (ERE) is shown as a green box.
B. Relative luciferase activity of the *IRX3* promoter reporter in response to co-expression with ERα in the presence or absence of estrogen (E2) or 4OH-tamoxifen (4OH-tam) in E2-deprived MCF-7 and MDA-231 cells. The data are derived from n=3 biological replicates from a single experiment. Bar graphs show means ±SD.
C. Genomic view of the IRX3 promoter and publicly available ChIP-seq tracks of ERα binding in MCF-7 cells in the absence (E2-) or presence of estrogen (E2+). Data derived from published studies (GSE32222 and GSE14664) (3,4). Indicated signal intensity is scaled according to positive ERα ChIP-seq signals in other loci.

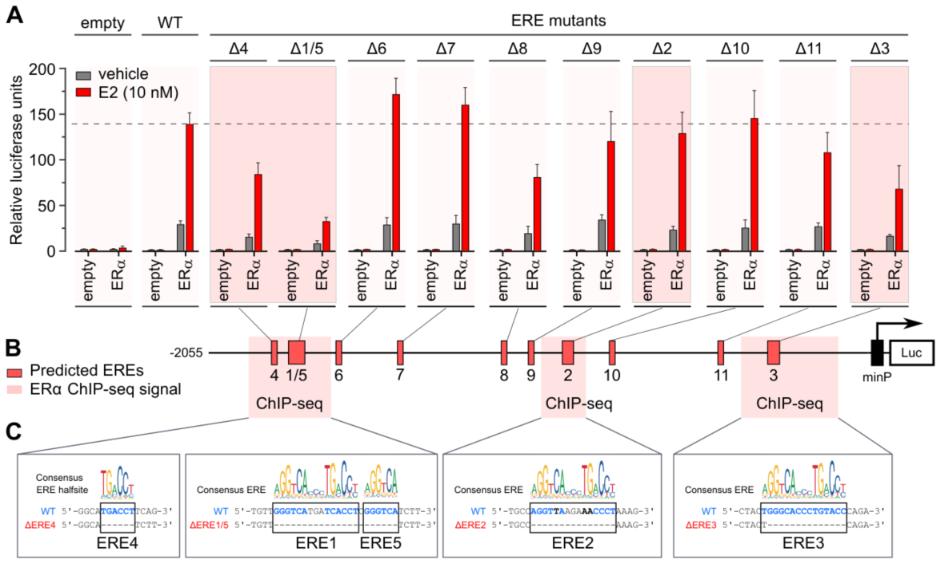

**Figure S3. The presence of several EREs at the enhancer region upstream of *IRX3* gene denotes ERα binding**

The WT full-length *IRX3* enhancer was cloned into the pGL4.23-minP-empty luciferase vector. Site-directed deletion of JASPAR-predicted EREs and half sites was performed and the effect on luciferase activity in response to overexpression of ERα and E2 treatment for 24 h in E2-deprived MDA-MB-231 cells was assessed.

A. Relative luciferase activity of each reporter construct. The data are derived from n=3 biological replicates and representative of three independent experiments. Bar graphs show means ±SD. Each mutated ERE is denoted as “Δ” followed by the ERE number.
B. Schematic illustration of the position of each ERE as dark red boxes. Regions overlapping with ERα ChIP-seq peaks in MCF-7 cells shown in light red boxes.
C. The consensus ERE shown as position weight matrix logo aligned with the genomic sequence of selected EREs of the enhancer.

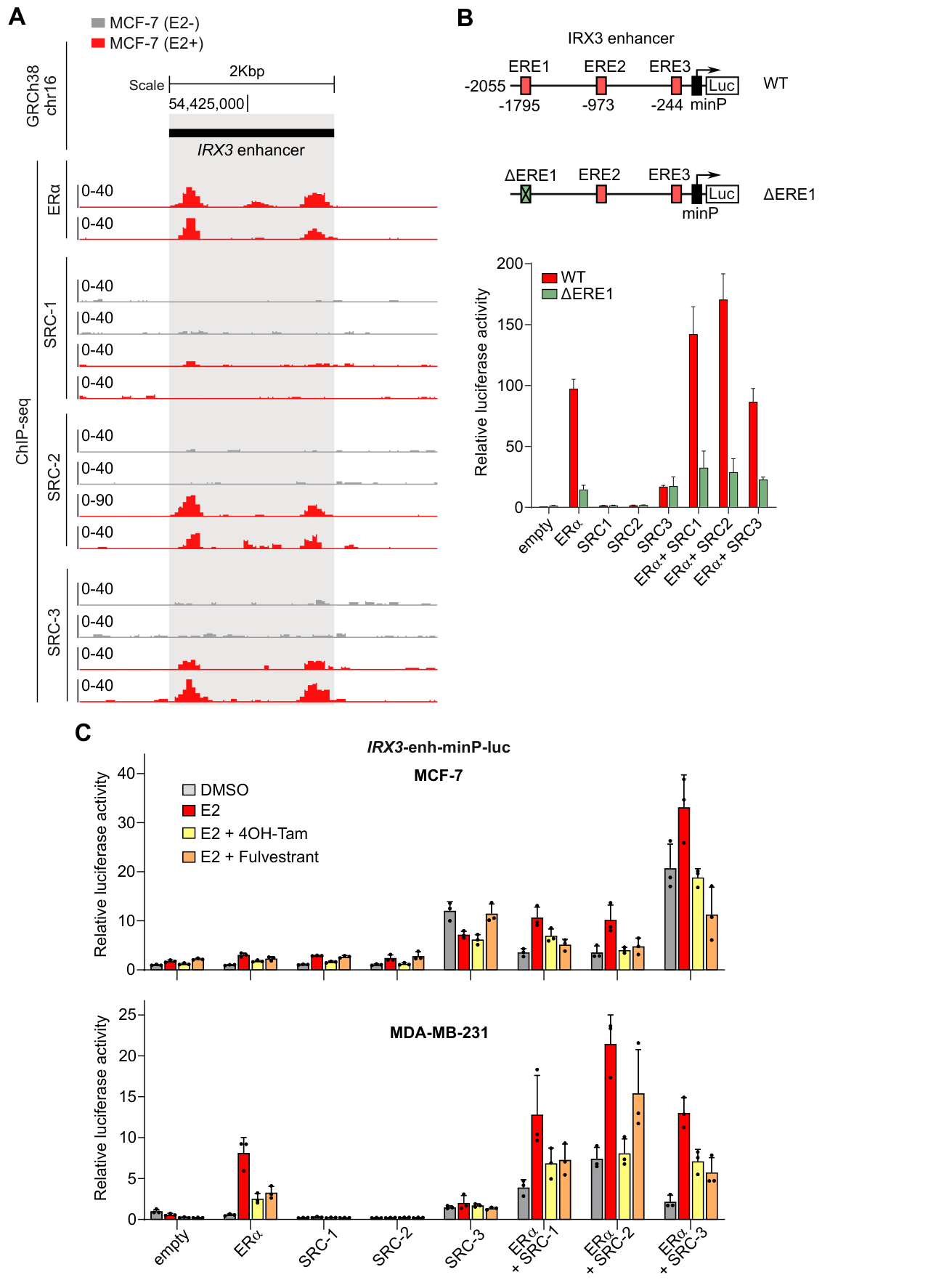

**Figure S4: IRX3 enhancer is activated by ERα coactivators**

A. ChIP-seq tracks showing genomic binding of ERα and steroid receptor coactivators (SRCs) to the *IRX3* enhancer in MCF-7 cells. The IRX3 enhancer is marked by a black bar and a grey box. The data are derived from a published study(5) (E-MTAB-785).
B. Luciferase reporter assay in MDA-MB-231 cells with overexpression of SRCs. Coactivators were overexpressed with or withoug ERα together with the IRX3 enhancer luciferase reporters in cells that were E2 deprived for 72h prior to treatment with E2 for another 24h followed by assessment of luciferase activity. The effect on the WT (red) and ΔERE1 (green) enhancers are shown. Data is derived from n=3 biological replicates from a single experiment. Bar graphs show means±SD.
C. Effects of 4OH-Tam and Fulvestrant on ER+ MCF-7 and ER- MDA-MB-231 cells. Steroid receptor co-activators (SRCs) were overexpressed with or without ERα overexpression together with the IRX3 enhancer luciferase reporter. The cells were subjected to E2 deprivation for 72h prior to treatment with vehicle, E2, 4OH-tamoxifen (4OH-tam) Tam) or Fulvestrant for 24h before assessment of the luciferase activity. Data are derived from n=3 biological replicates from a single experiment. Bar graphs show means ±SD.

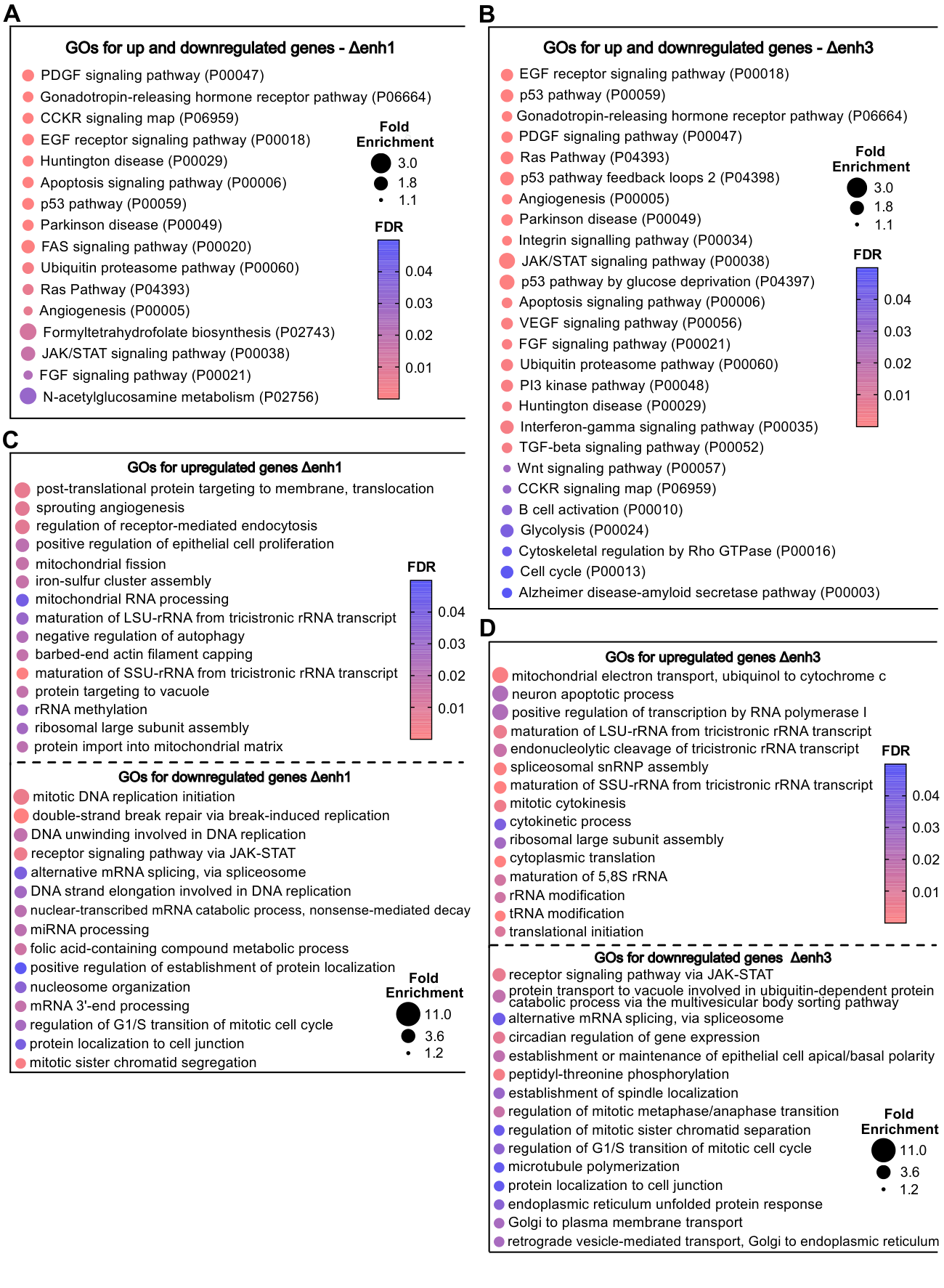

**Figure S5. Gene ontologies for differentially expressed genes in Δenhancer cells**

**A-B**. Differences in global gene expression between control and the Δenhancer clones with the strongest reduction in IRX3 protein levels were assessed by RNA-seq, with n $=$ 4 biological replicates in a single experiment. All significant PANTHER pathways (FDR $<$ 0.05) of differentially expressed genes (*p_adj_* $<$ 0.01) are shown.

**C-D.** The effect of IRX3 knockdown by genomic deletion of the upstream enhancer on global gene expression was assessed by RNA-seq (n=4 biological replicates from a single RNA-seq run). Panther GO – Slim gene ontology analyses (FDR <0.05) was performed on up- and down-regulated DEGs (*p_adj_* < 0.01) for C, Δenh1 and D, Δenh3. Δenh, Δenhancer*.* Top 15 enriched gene ontology (GO) categories are shown.

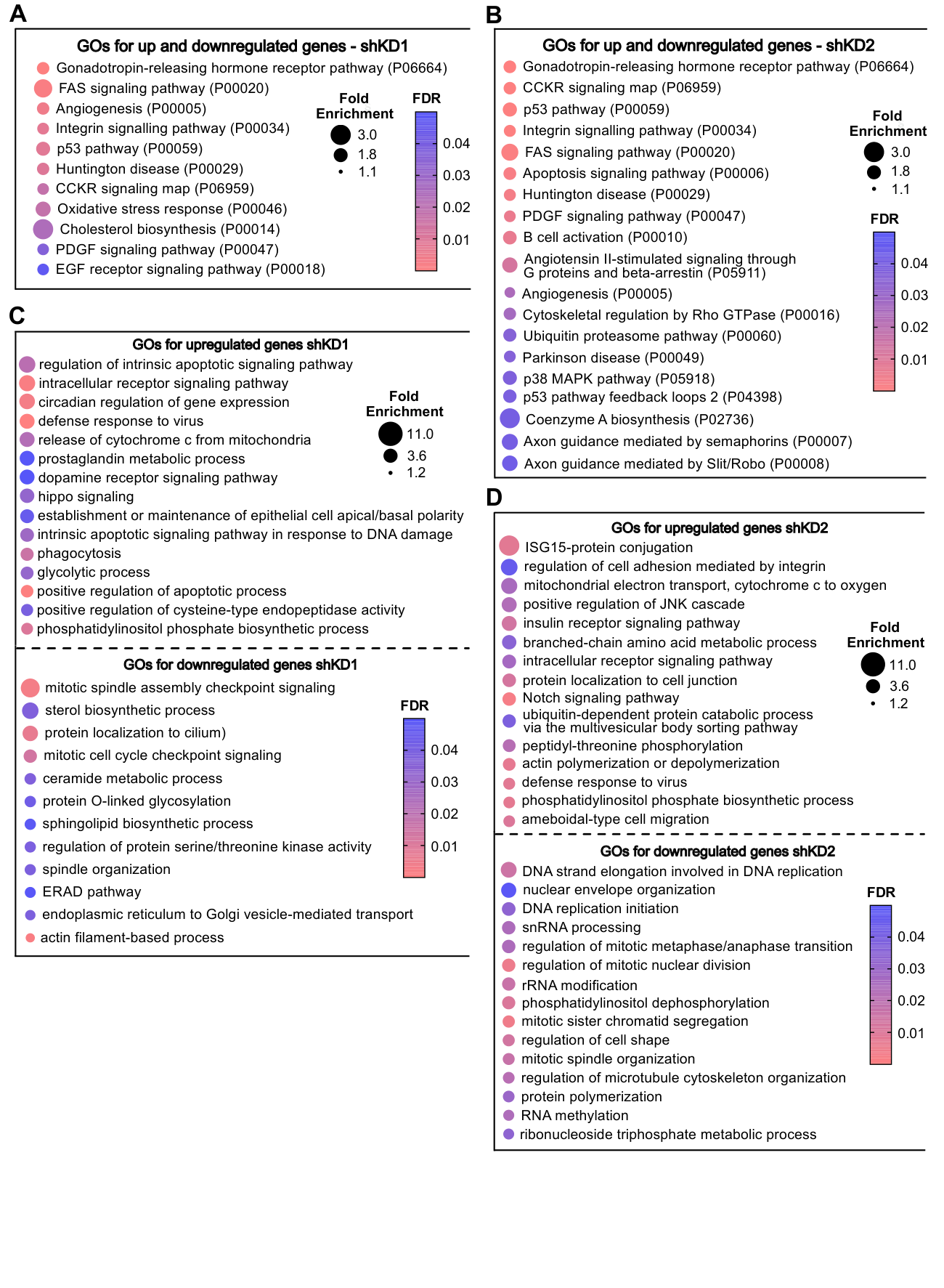

**Figure S6. Gene ontologies for differentially expressed genes in shIRX3 cells**

**A-B**. Differences in global gene expression between control and shKD cells were assessed by RNA-seq, with n $=$4 biological replicates in a single experiment. Panther pathways GO analyses of all DEGs (*p_adj_* $<$0.01) are shown (**Supplementary Dataset 1**).

**C-D.** The effect of IRX3 knockdown directly using targeting shRNAs on global gene expression was assessed by RNA-seq (n=4 biological replicates from a single RNA-seq run). Panther GO – Slim gene ontology analyses (FDR <0.05) was performed on up- and down-regulated DEGs (*p_adj_* < 0.01) for C, shKD1 and D, shKD2 (**Supplementary Dataset 1**). shKD, shRNA targeting *IRX3.* Top 15 enriched gene ontology (GO) categories are shown.

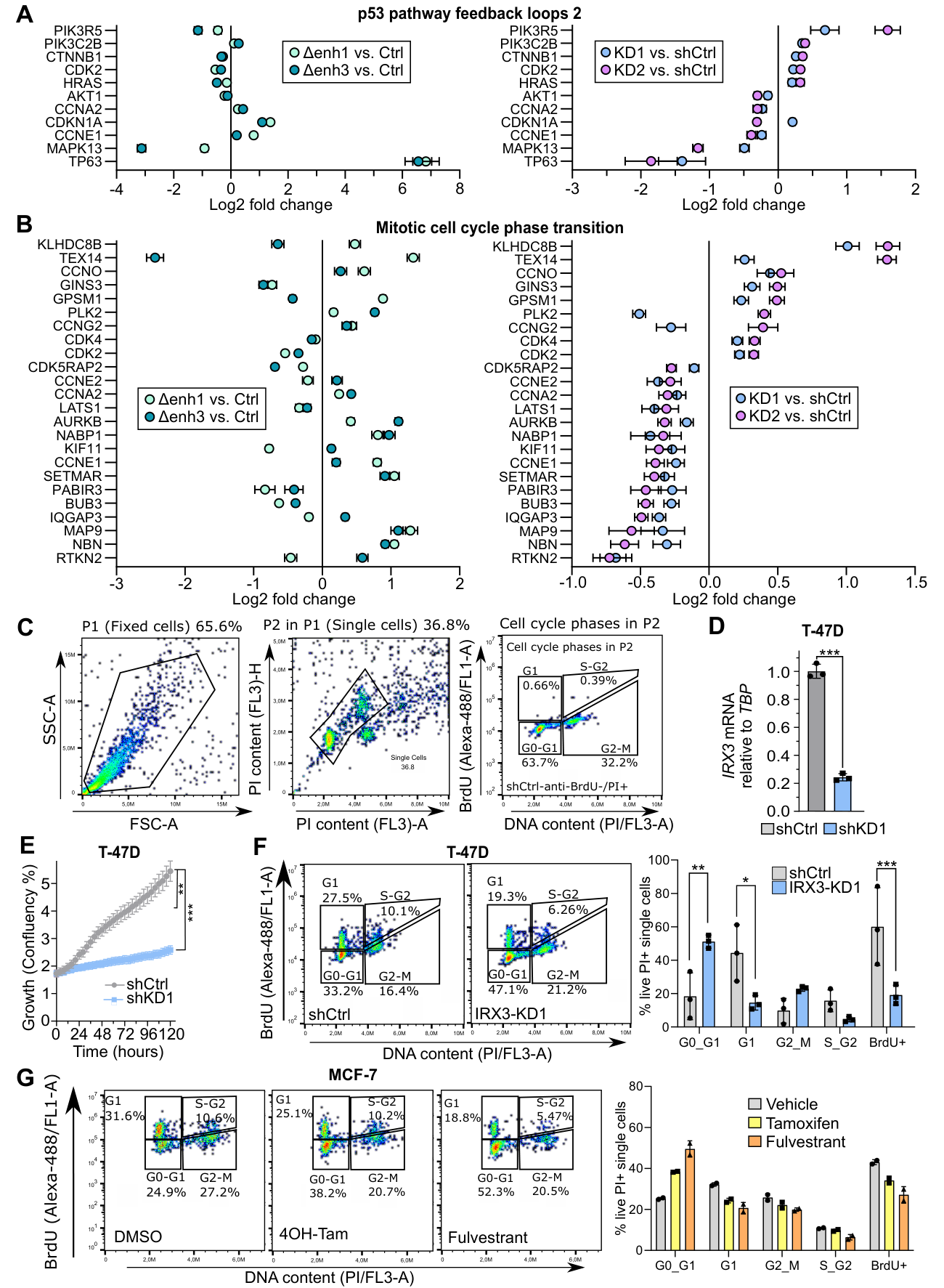

**Figure S7. Decreased IRX3 levels promotes cell cycle arrest and reduced proliferation**

**A-B.** Fold changes of differentially expressed genes comparing MCF-7 Δenhancer vs. control (left panel) and *IRX3-*KD vs. shCtrl cells (right panel) comprising the GO category “p53 pathway feedback loops” (A) and “Mitotic cell cycle process” (B) (**Supplementary Dataset 1**). Data are derived from n = 4 biological replicates in a single RNA-seq experiment. Graphs show mean log_2_FC ±SE.
**C.** Gating strategies in the two-dimensional (2D) cell cycle analysis. The total cell population was pre-gated with FSC-A vs. SSC-A (left panel) and FL3-A vs. FL3-H (middle panel) to eliminate cell debris and cell doublets, respectively. The BrdU against DNA content was defined on the pre-gated cells (right panel, an example of negative ctrl that was stained with anti-IgG instead of anti-BrdU antibody). The gates represent BrdU-negative cells in G0-G1, G2-M and BrdU-positive G1 and S-G2 (Cells which have undergone at least one round of cell division since incorporation of BrdU).
**D.** The expression of *IRX3* in ER+ T-47D control and *IRX3*-KD cells (shKD1) was measured by qPCR, normalized to *TBP*. ****p_adj_* 0.001; Student’s two-sided t-test. Data are derived from n=3 biological replicates from a single experiment. Bar graphs show means ± SD.
**E.** Growth curves of T-47D control and *IRX3-*KD cells as determined by the IncuCyte S3 Live-Cell system and is presented as percentage of confluence normalized to *t*=0. Data are derived from n=8 biological replicates from a single experiment. ***p_adj_* < 0.01, ****p_adj_* <0.001; Two-Way Anova with Holm-Sidak correction for multiple testing. Graphs show means ±SD.
**F.** 2D cell cycle assay. Asynchronous T-47D control and *IRX3*-KD cells were pulse-labeled with BrdU for 1 h and left to grow overnight in BrdU-free medium. Harvested cells were fixed and stained with Alexa-488 conjugated anti-BrdU and propidium iodide (PI). Bivariate contour plots of PI vs BrdU from representative samples are shown with gates for G1, S-G2, G2-M and G0-G1 (left panel). Percentage of cells in each gate is shown in the bar plot (right panel). Data are derived from n=3 biological replicates from a single experiment. **p_adj_* 0.05, ***p_adj_* 0.01, ****p_adj_* 0.001; Multiple two-sided Student’s t-tests with Holm-Sidak correction for multiple testing. Bar graphs show means ±SD.
**G.** 2D cell cycle analysis of ER+ MCF-7 treated with either 250nM 4OH-Tam or 100nM Fulvestrant. Cells were stained and analyzed as in (F). Bivariate contour plots of PI vs BrdU from representative samples are shown with gates for G1, S-G2, G2-M and G0-G1 (left panel). Percentage of cells in each gate is shown in bar plots (right panel), n=2 biological replicates from two individual experiments. Bar graphs show means ±SD.

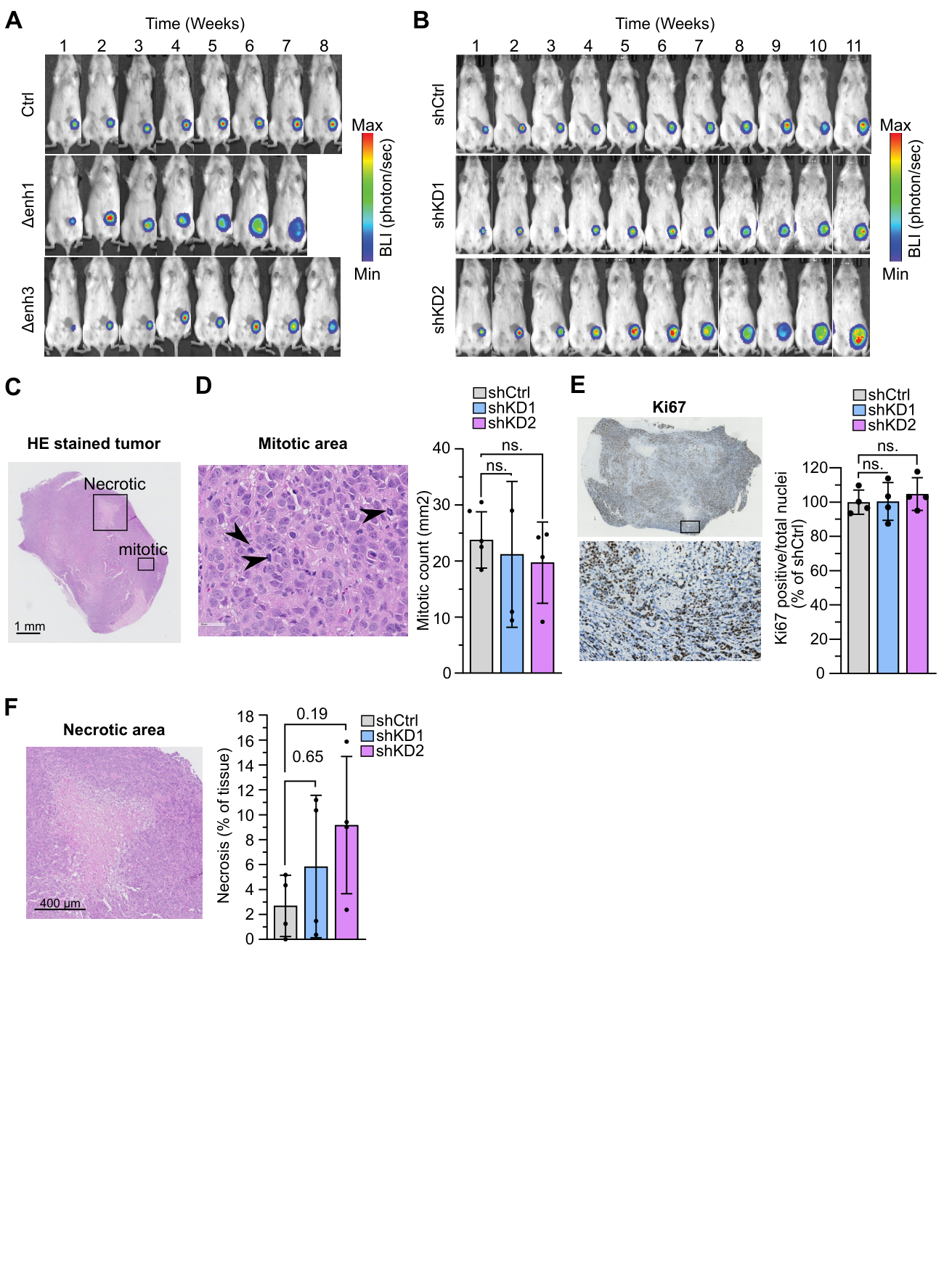

**Figure S8. IRX3 depletion in ER+ breast cancer cells increases *in vivo* tumor growth and necrosis**

A. *In vivo* bioluminescence images of representative mice with implanted control and ∆enhancer cells from week 1-8 post implantation (one representative mouse per implanted cell line is shown). ∆enh1-recipient

mice were sacrificed on week 7 post-implantation due to the large primary tumor size.

B. *In vivo* bioluminescence images of mice with implanted control and MCF-7 shIRX3 knockdown cells from week 1-11 post implantation (one representative mouse per implanted cell line is shown).
C. Representative HE-stained section of the primary tumor of a shCtrl-recipient mouse, in which necrotic areas and tumor cell mitoses are present.
D. Scoring the mitotic cells per mm^2^ in mitotic hotspots from HE-stained section prepared from the excised primary tumors from **Fig. 5G**, n=4. Data is shown as means $\pm$ SD, Non-parametric Kruskal-Wallis test was performed with Dunnett’s multiple comparison test.
E. Quantified Ki-67 positive nuclei in FFPE tumor sections prepared from the excised primary tumors from **Fig. 5G**, n=4. Immunohistochemical staining was performed using a monoclonal mouse anti-human Ki67 antibody. Data shown as means $\pm$ SD. Ordinary one-way Anova was done with Dunnett’s multiple comparison test.
F. Quantified necrotic area in HE-stained section prepared from the excised primary tumors from **Fig. 5G**, n =4. Data shown as means $\pm$ SD. Non-parametric Kruskal-Wallis test was done with Dunnett’s multiple comparison test.

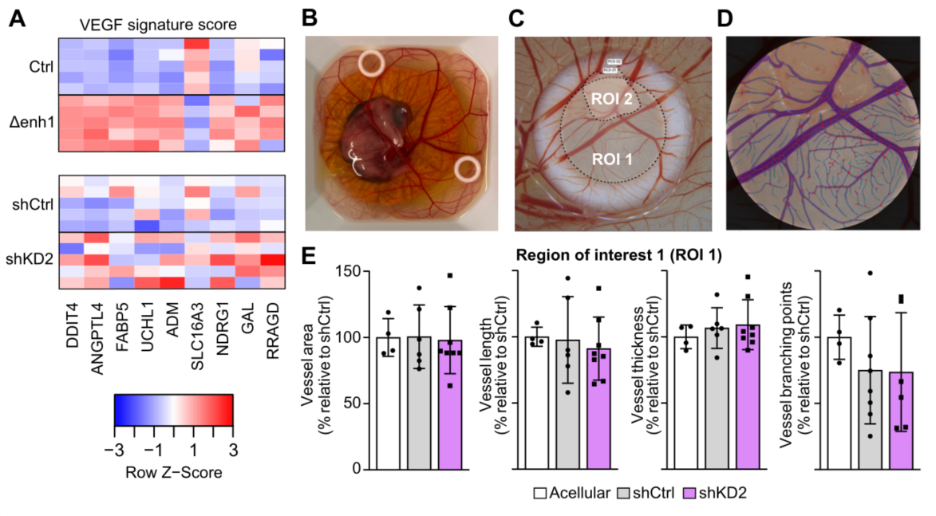
**Figure S9. Reduced IRX3 levels promote vascularization *in vivo***

A. RNA-sequencing of mouse primary tumors (**Fig. 5**) were performed. Heatmap presents the z-score of each gene expression within the VEGF gene signature in ctrl, Δenh1, shCtrl and shKD2 (n =5 in each group).
B. A representative image of an *ex ovo* chicken embryo at day 12 post fertilization, where shCtrl and shKD2 MCF-7 cells were prepared in Matrigel and implanted in Teflon rings on top of the chorioallantoic membrane (CAM) of chicken embryos *ex ovo* at embryonic day 7. After 5 days, Teflon rings with the CAMs containing the MCF-7 cells were dissected and imaged, before quantifying blood vessels by stereomicroscopy and deep-learning imaging analysis.
C. A representative picture of a Teflon ring on the CAM of a chicken embryo on embryonic day 12, containing two different regions of interest (ROI), where ROI2 is the MCF-7 cells in Matrigel that has formed a spheroid and ROI 1 is the surrounding CAM area inside the Teflon ring.
D. A microscopic image showing deep-learning imaging analysis of the chicken embryo CAM blood vessels inside the spheroid and surrounding CAM area.
E. Quantification of blood vessels within ROI 1 based on deep-learning imaging analysis. Data are derived from n=4-7 independent implants in a single experiment. Acellular is Teflon rings with only Matrigel, but no implanted cells. Data shown as means ± SD.

**References**

1.	Pal B, Chen Y, Vaillant F, Capaldo BD, Joyce R, Song X*, et al.* A single-cell RNA expression atlas of normal, preneoplastic and tumorigenic states in the human breast. EMBO J **2021**;40:e107333

2.	Helland T, Gjerde J, Dankel S, Fenne IS, Skartveit L, Drangevag A*, et al.* The active tamoxifen metabolite endoxifen (4OHNDtam) strongly down-regulates cytokeratin 6 (CK6) in MCF-7 breast cancer cells. PLoS One **2015**;10:e0122339

3.	Welboren WJ, van Driel MA, Janssen-Megens EM, van Heeringen SJ, Sweep FC, Span PN*, et al.* ChIP-Seq of ERalpha and RNA polymerase II defines genes differentially responding to ligands. EMBO J **2009**;28:1418-28

4.	Ross-Innes CS, Stark R, Teschendorff AE, Holmes KA, Ali HR, Dunning MJ*, et al.* Differential oestrogen receptor binding is associated with clinical outcome in breast cancer. Nature **2012**;481:389-93

5.	Zwart W, Theodorou V, Kok M, Canisius S, Linn S, Carroll JS. Oestrogen receptor-co-factor-chromatin specificity in the transcriptional regulation of breast cancer. EMBO J **2011**;30:4764-76
